# *In planta*-specific transcriptional regulatory circuit regulates expression of *MoHTR1*, a nuclear effector gene of *Magnaporthe oryzae*

**DOI:** 10.64898/2026.03.02.709172

**Authors:** Yoon-Ju Yoon, Hyunjun Lee, Seongbeom Kim, Hyunjung Chung, Chang Hyun Khang, You-Jin Lim, Yong-Hwan Lee

## Abstract

During host-pathogen interactions, fungal pathogens secrete effector proteins into host cells to manipulate the host immune system and facilitate infection. Although many effector genes are highly expressed during infection stages, there is limited information on the mechanisms regulating their *in planta* expression. Here, we characterize the promoter of *MoHTR1*, a nuclear effector gene of the rice blast fungal pathogen, to elucidate its *in planta*-specific expression. Using promoter deletion and mutation analyses, we identified a core *cis*-element (TATTTCGT) within the *MoHTR1* promoter, designated the *in planta* active (IPA) element, which is crucial for *in planta*-specific expression. The IPA element is responsible for the expression of not only *MoHTR1*, but also other effector genes including a known effector *Slp1*. Furthermore, the IPA element enables the *in planta* expression of *MobZIP14*, a gene specifically expressed during vegetative growth. The IPA element plays a critical role in fungal virulence by enabling *MoHTR1* expression and regulating host immune responses. Bioinformatic and DNA-protein interaction analyses revealed that RGS1, a transcription factor containing a winged-helix binding domain, acts as a transcriptional regulator of *MoHTR1* by directly binding to the IPA element. Our findings provide new insights into the regulatory mechanisms driving the *in planta*-specific expression of fungal effector genes.

## INTRODUCTION

Among the various factors that induce plant disease, effectors are crucial virulence determinants during plant-pathogen interactions (Kunkel and Chen, 2006; Pontes et al., 2020). Plant pathogens secrete effectors to suppress the plant immune system or modulate host metabolism for successful infection (Giraldo and Valent, 2013). Effectors are categorized into three classes based on their distinct localizations during infection: apoplastic, cytoplasmic, and nuclear effectors. Apoplastic effectors accumulate in the space between invasive hyphae (IH) and the extra-invasive hyphal membrane (EIHM), while cytoplasmic and nuclear effectors are translocated into the host cell via the biotrophic interfacial complex (BIC) (Giraldo and Valent, 2013). Nuclear effectors are transported into the nucleus to modulate the expression of host immunity-associated genes (Kim et al., 2020a). Plant pathogens coordinate the expression of these diverse effector genes to evade the host immunity and facilitate infection. Many effector genes are highly induced transcriptionally during infection stages but show very low or no expression in the absence of the host plant (Yan et al., 2023). However, the mechanisms regulating effector gene expression during the plant-pathogen interactions remains largely unexplored.

Promoters are key controllers of gene expression and can be classified as constitutive or inducible based on their expression patterns. Both constitutive and inducible promoters contain *cis*-elements, which are the sequences bound by transcription factors to regulate gene expression. In contrast to constitutive promoters, inducible promoters are activated by specific stimuli (Hernandez-Garcia and Finer, 2014; Villao-Uzho et al., 2023). Understanding the regulatory networks of gene expression is crucial for interpreting the complexities of cellular functions, developmental processes, and responses to various stresses (Sijacic et al., 2018; Wang et al., 2015). Several transcriptional regulatory mechanisms have been identified in plants, particularly those involving *cis*-element and transcription factors related to abiotic and biotic stresses. For instance, W-box and GCC-box are well-known pathogen-inducible *cis*-elements in rice and other plants (Kong et al., 2018; Van der Does et al., 2013). The W-box is directly bound by WRKY transcription factors to regulate expression of defense-related genes (Ishihama et al., 2011; Xiao et al., 2013). In bacterial plant pathogens, the plant-inducible promoter (PIP) box is a well-known *cis*-element in *Xanthomonas* and *Ralstonia* species (Cunnac et al., 2004). The PIP box is conserved in the promoter regions of hypersensitive response and pathogenicity (*hrp)* genes and some type III secreted (T3S)-effector genes, which are specifically expressed during infection and are regulated by the HrpX transcription factor (Jiang et al., 2009). However, compared to host plant and bacterial pathogens, the mechanisms regulating *in planta*-specific expression of effector genes in plant pathogenic fungi remains poorly understood.

Rice blast, caused by *Magnaporthe oryzae*, is one of the most devastating diseases affecting rice and wheat, leading to annual economic losses that could feed 60 million people (Nalley et al., 2016). Due to its socio-economic importance and genetic tractability, *M. oryzae* has been used as a model organism for studying the interactions between plants and pathogens. The rice blast pathogen undergoes various developmental processes to invade host cells, including conidial adhesion, conidial germination, appressorium formation, penetration into plant cells, and the development of invasive hyphae (Wilson and Talbot, 2009). Numerous effectors have been studied in the rice blast pathogen, including BAS4 and Slp1, typical apoplastic effectors, and PWL2 and Avr-Pita, cytoplasmic effectors (Abbas et al., 2022; Lo Presti et al., 2015; Oliveira-Garcia et al., 2024). Recently, MoHTR1 and MoHTR2, the nuclear effectors, were reported to be secreted and translocated into host nuclei to manipulate host immunity through transcriptional reprogramming (Kim et al., 2020a). Among many effectors reported in *M. oryzae*, only a few studies have identified the *cis*-elements and transcription factors involved in effector gene expression. For example, a 12 bp (TTATGCAAGCTT) sequence within the *PWL2* promoter acts as a *cis*-element for biotrophy-specific expression (Zhu et al., 2021). MoEITF1 and MoEITF2 are transcription factors that regulate effector gene expression during early infection stages, and RGS1 has been identified as a regulator of effector gene expression during conidial stage (Cao et al., 2022; Tang et al., 2023). However, no studies have simultaneously identified both the transcription factors and the corresponding *cis*-elements to understand *in planta*-specific expression of effector genes, leaving the transcriptional network of effector gene expression largely unexplored.

In this study, we characterized the promoter of *MoHTR1*, the nuclear effector gene, to decipher it’s *in planta*-specific expression. We identified an 8-bp *in planta* active (IPA) element of *MoHTR1*, which is essential for the *in planta*-specific expression of not only *MoHTR1* but also other effector genes including *Slp1*, an apoplastic effector gene. The IPA element also enables the *in planta* expression of *MobZIP14*, a gene specifically expressed during vegetative growth. The IPA element of *MoHTR1* plays crucial roles in fungal virulence and regulating host immune response. Furthermore, we demonstrated that the transcription factor RGS1 directly binds to the IPA element and regulates *in planta*-specific expression of *MoHTR1*. These findings provide new insights into the regulatory mechanisms of *in planta*-specific expression of fungal effector genes.

## RESULTS

### *MoHTR1* is specifically expressed during *in planta* stages

To determine the expression patterns of *MoHTR1* during rice infection, we performed qRT-PCR using RNA samples from *M. oryzae*-infected rice leaves at 24, 48, and 72 hours post inoculation (hpi), with mycelial RNA as a control. *MoHTR1* expression was induced and increased significantly during infection, reaching at least 600-fold at 24 hpi compared to the mycelial growth stage, peaking at a remarkable 2,300-fold at 48 hpi, and then reducing to 660-fold at 72 hpi **(Supplemental Fig. 1A)**. To further validate the *in planta*-specific expression of *MoHTR1* at single-cell resolution, we used *M. oryzae* strain carrying a *MoHTR1* promoter fused with GFP at various developmental and infection stages. Strong GFP fluorescence was observed in invasive hyphae (IH), while no fluorescence signal was detected in mycelia, conidia, germinated conidia, or appressoria **(Supplemental Fig. 1B)**, consistent with qRT-PCR results **(Supplemental Fig. 1A)**.

To investigate whether the *in planta*-specific expression of *MoHTR1* occurs consistently across different host and non-host plants and whether it is induced only in living cells, we inoculated a fungal transformant co-expressing EF1αpro:mRFP and MoHTR1pro:sGFP into living and heat-killed (70 ℃) cells of rice, barley (another host for *M. oryzae*), and onion (a non-host plant for *M. oryzae*). GFP fluorescence was clearly detected in IH within living rice, barley, and onion cells, but was absent in heat-killed cells **(Supplemental Fig. 2A)**. Furthermore, while GFP signal remained detectable in rice sheath cells exposed to 42 ℃ heat stress, no signal was detected in cells killed by freezing **(Supplemental Fig. 2B and C)**.

To examine whether the *in planta*-specific expression of *MoHTR1* is regulated by immune signaling pathways, we treated infected rice sheath cells with inhibitors targeting each pathway including Ca^2+^, MAP kinase, SA, JA, and ROS. GFP fluorescence was consistently observed in IH across all inhibitor-treated samples **(Supplemental Fig. 3A and B)**. Furthermore, when we applied cutin monomers in heat-killed cells, none of the treatments restored *MoHTR1* promoter activity **(Supplemental Fig. 3C)**.

### The 8-bp *in planta* active (IPA) element is essential for *in planta*-specific expression of MoHTR1

To identify the *cis*-acting element regulating *in planta*-specific expression of *MoHTR1*, which we termed the *in planta* active (IPA) element, we generated transcriptional reporter constructs by fusing various truncated *MoHTR1* promoters with sGFP (superfolder green fluorescent protein). Each construct was transformed into *M. oryzae*, and the resulting transformants were inoculated into rice sheaths to observe sGFP expression using a fluorescence microscope. Our initial set of truncated promoters, designed with 150-bp sequential deletions between 900 bp and 300 bp upstream of the *MoHTR1* translation start site, showed strong GFP fluorescence for promoters longer than 450 bp (*MoHTR1_pro450_*), but no GFP fluorescence was observed with the 300-bp promoter (*MoHTR1_pro300_*) **(Supplemental Fig. 4A)**. Further successive 50-bp deletions between *MoHTR1_pro450_* and *MoHTR1_pro300_* revealed no GFP fluorescence with the 400-bp construct (*MoHTR1_pro400_)* or shorter, indicating that the IPA element of *MoHTR1* is located within 400-450 bp upstream of the *MoHTR1* translation start site **(Supplemental Fig. 4B)**. To identify the core sequence of the IPA element, we subdivided the putative IPA element region into 25-bp segments **(Fig. 1A)**. The region from -400 to -425 bp upstream of *MoHTR1* translation start site was critical for *in planta*-specific expression **(Fig. 1B)**. Given that DNA elements regulating transcription are typically about 10 bp long (Stewart et al., 2012), we further divided the target IPA element region into three motifs **(Fig. 1C)**. Each motif was designed to overlap by 3-4 bp (IPA motif 1: -425 to -415, IPA motif 2: -419 to -409, IPA motif 3: -412 to -401 relative to the translation start site of *MoHTR1*) and was mutated using transversion substitutions, where purine bases (A, G) were replaced with pyrimidine bases (C, T), or vice versa. We found no GFP fluorescence for *MoHTR1_IPA1_mut_*, while *MoHTR1_IPA2_mut_* and *MoHTR1_IPA3_mut_* exhibited strong GFP signals comparable to those of the wild-type construct **(Fig. 1D)**.

**Figure 1.**
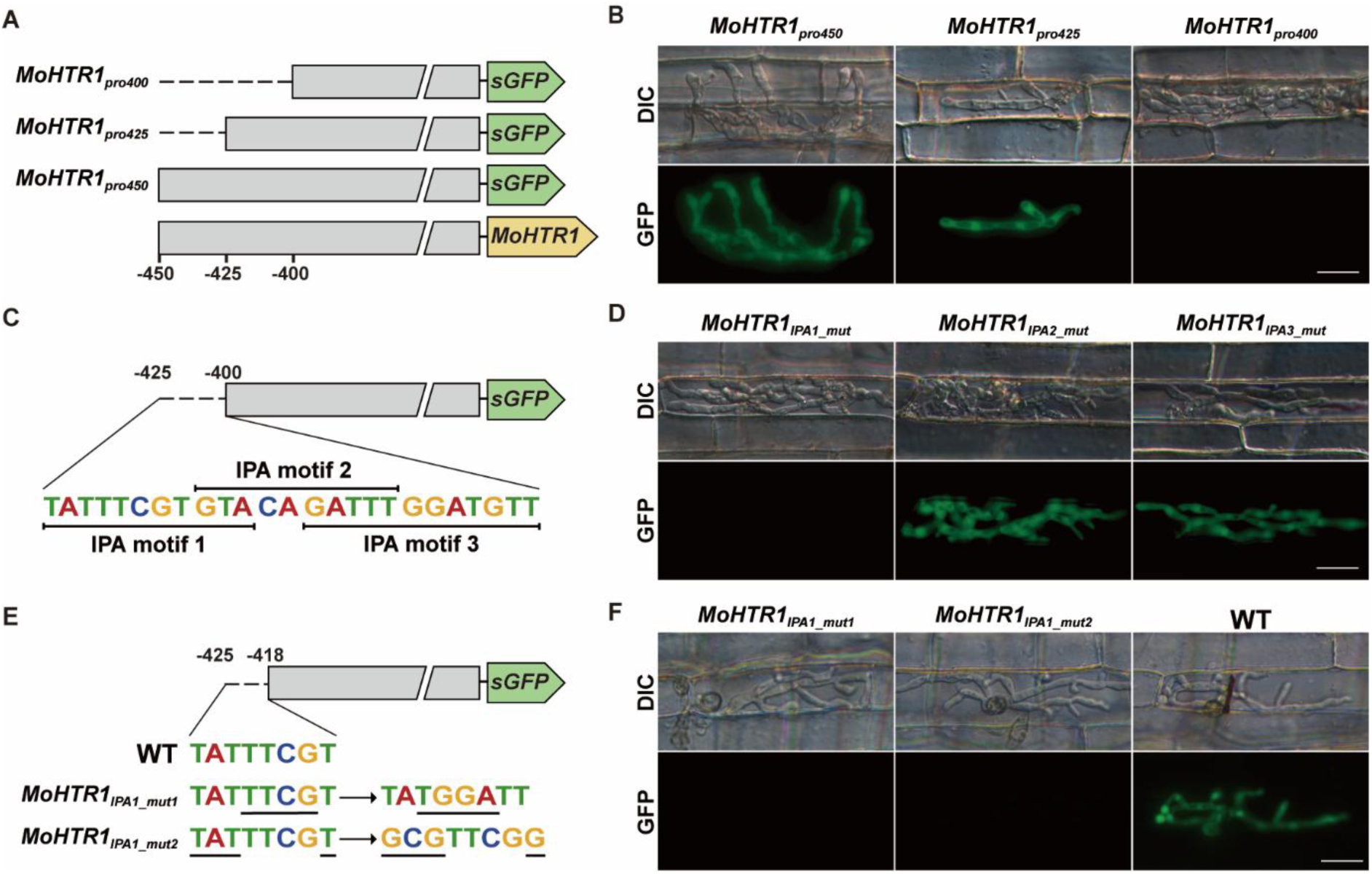
The 8-bp sequence of IPA element is required for *in planta*-specific expression of *MoHTR1*. **(A)** Schematic representations of truncated *MoHTR1* promoter fused with GFP to determine the minimal region required to drive GFP expression *in planta*. **(B)** GFP fluorescence of various *MoHTR1* promoter constructs in the invasive hyphae. Each strain was inoculated in rice sheath cells for 32 h. Scale bar indicates 20 μm. **(C)** Schematic diagram of the putative IPA element region of *MoHTR1*. The 25-bp of target IPA element region is divided into three 12-bp parts and defined as IPA motif1 (-425 to -415 bp), IPA motif2 (-419 to -409 bp), and IPA motif3 (-412 to -401 bp). **(D)** GFP fluorescence of the IPA motif mutation constructs in the invasive hyphae. Each strain was inoculated in rice sheath cells for 32 h. Scale bar indicates 20 μm. **(E)** Schematic diagram of *MoHTR1* promoter constructs with targeted mutations in the IPA element. Two constructs were generated: *MoHTR1_IPA1_mut1_*, with a transversion mutation in the PIP box-like sequence within the IPA element, and *MoHTR1_IPA1_mut2_*, with a mutation in the remaining region excluding the PIP box-like sequence. Both constructs were fused to sGFP to evaluate promoter activity during infection. **(F)** GFP fluorescence of *MoHTR1* IPA element mutants in the invasive hyphae. Each strain was inoculated in rice sheath cells for 32 h. Scale bar indicates 20 μm.

To further validate promoter activity, we measured the GFP intensity in IH at 30 hpi for five strains (*MoHTR1_pro400_, MoHTR1_IPA1_mut_, MoHTR1_IPA2_mut_, MoHTR1_IPA3_mut_,* and *MoHTR1_pro425_*). Promoter activity was normalized on a scale from 0 to 1, where 0 indicates no fluorescence, and 1 represents the average intensity observed in IH for the positive control, *MoHTR1_pro425_*. The GFP intensities for *MoHTR1_IPA2_mut_* and *MoHTR1_IPA3_mut_* were 0.9 and 0.86, respectively, showing no significant difference from the positive control, *MoHTR1_pro425_*. In contrast, *MoHTR1_IPA1_mut_* and the negative control, *MoHTR1_pro400_*, showed intensities near 0 **(Supplemental Fig. 4C),** indicating the loss of promoter activity.

The PIP box (TTCG-N_16_-TTCG) is a well-known *cis*-element involved in *in planta* expression in bacterial pathogens (Cunnac et al., 2004). To examine whether the PIP box-like sequence within the IPA element contributes to its function, we generated two mutants: *MoHTR1_IPA1_mut1_* (TAT<u>TTCG</u>T), in which the PIP box-like region within the IPA element was transversion-mutated, and *MoHTR1_IPA1_mut2_* (<u>TAT</u>TTCG<u>T</u>), in which the remaining region of the IPA element excluding the PIP box-like sequence was mutated **(Fig. 1E)**. GFP intensity of *MoHTR1_IPA1_mut1_* was significantly reduced (0.22) compared to the positive control strain, *MoHTR1_pro425_*, while *MoHTR1_IPA1__mut_2_* showed almost no detectable GFP signal **(Fig. 1F and Supplemental Fig. 4D)**. These results demonstrate that the entire 8-bp (TATTTCGT) sequence of the IPA element is essential for *in planta*-specific expression of *MoHTR1*, and that this element is functionally distinct from the bacterial PIP box. To further determine whether reduced promoter activity caused by mutations in the IPA element also affects subcellular localization of MoHTR1 during infection, we generated fungal transformants expressing MoHTR1 fused to mRFP (MoHTR1:mRFP) under the native promoter and under promoters carrying mutations in the IPA element. In the native promoter strain, MoHTR1:mRFP accumulated in BICs and in rice nuclei, whereas no detectable mRFP fluorescence was observed at BICs or in host nuclei in either strain harboring IPA element mutations **(Supplemental Fig. 4E)**.

The *MoHTR1* IPA element is crucial for the *in planta*-specific expression of other effector genes in *M. oryzae*.

Our bioinformatic analysis revealed that 415 out of the 12,593 genes in the *M. oryzae* genome contain the *MoHTR1* IPA element sequence within 1 kb upstream of their start codon **(Fig. 2A)**. Among these, 234 genes (56.4 %) were identified as up-regulated during infection, 110 genes showed no significant change in expression, and 71 genes were down-regulated compared to mycelial stage based on *in planta* RNA-seq data (Jeon et al., 2020) **(Fig. 2A and Supplemental Data 1)**. These 234 genes include two previously characterized effector and pathogenicity gene, *Slp1* and *MoHDA1*, respectively, and four effector candidate genes (Kim et al., 2019; Kim et al., 2020b; Mentlak et al., 2012) . Our qRT-PCR analysis further confirmed that these six genes exhibited the highly induced expression in IH that is similar to *MoHTR1* **(Fig. 2B),** suggesting that a common regulating mechanism for *in planta* expression is shared by *MoHTR1* and other effector genes. Notably, the IPA element responsible for regulating the expression of the nuclear effector gene, *MoHTR1*, is also implicated in the expression of the apoplastic effector gene, *Slp1*, whose protein suppresses PAMP-triggered immunity (Mentlak et al., 2012). To confirm the role of the IPA element in the *in planta*-specific expression of effector genes, we mutated the IPA sequence in *Slp1* promoter. A comparison of GFP fluorescence intensity under the native promoter of *Slp1* (*Slp1_pro825_*) and the transversion mutated promoter of *Slp1* (*Slp1_IPA_mut_*) showed a strong GFP signal in the IH of *Slp1_pro825_* but not in *Slp1_IPA_mut_*. These results indicate that the IPA element is essential for the *in planta*-specific expression of effector genes, including the nuclear effector (*MoHTR1*) and the apoplastic effector (*Slp1)* **(Fig. 2C)**. It is worth noting that both genes modulate host immunity with their proteins targeting distinct host compartments, the host nuclei and the apoplast, respectively.

**Figure 2.**
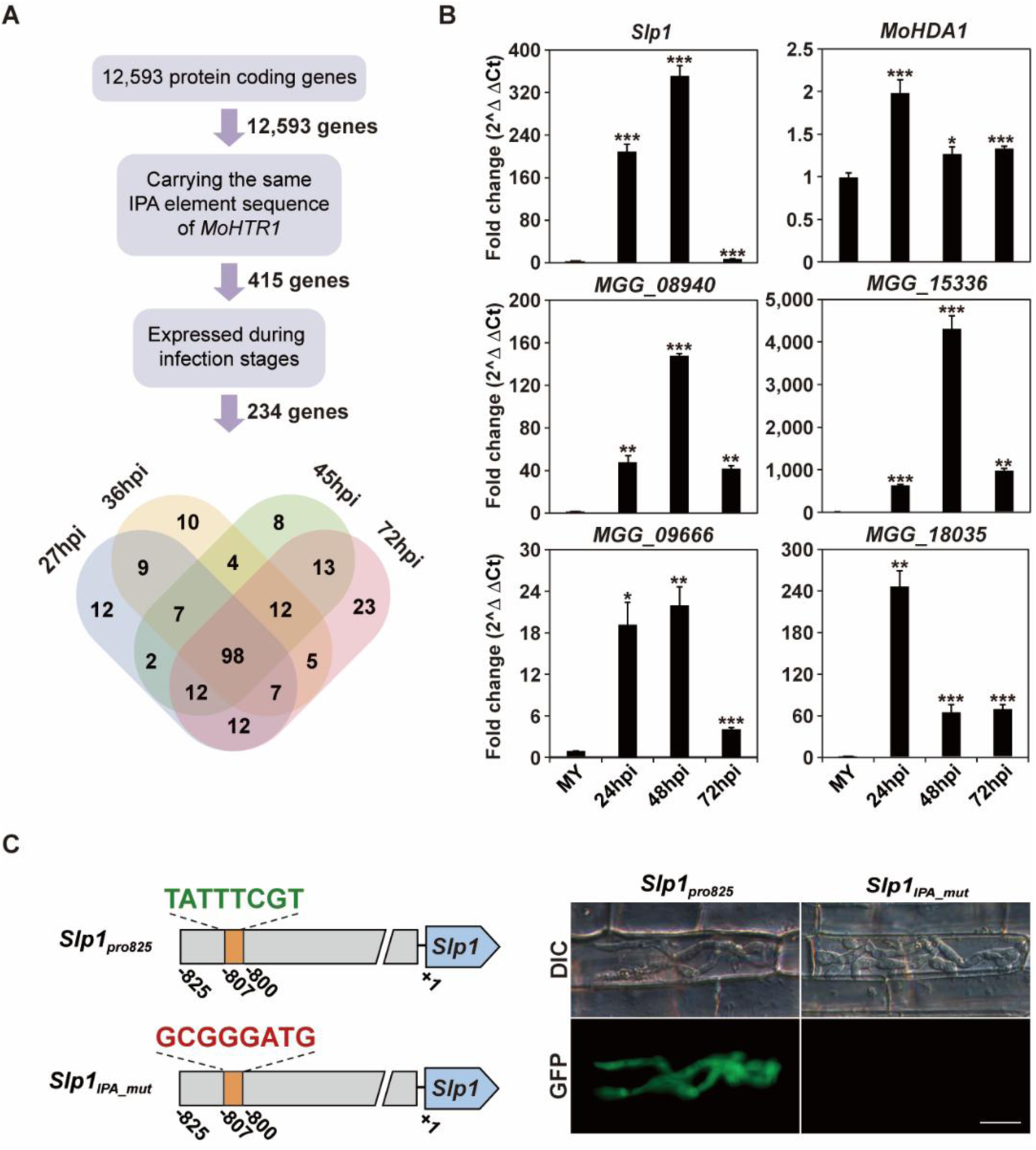
The IPA element of *MoHTR1* is crucial for regulation the expression of genes at the *in planta* stage. **(A)** Diagram of filtering pipeline for selecting the genes carrying the same *MoHTR1* IPA element sequence which highly expressed during the infection stages. See Supplemental Data 1 for promoter sequences, expression pattern at *in planta* stage, and the presence/absence of a signal peptide. **(B)** qRT-PCR analysis to verify the expression patterns of selected genes, which contain the same IPA element sequence of *MoHTR1*. Relative expression at all infection stages was calculated using the 2^−ΔΔCt^ method with mycelial stage as a reference, and *β-tubulin* was used for normalization. Mean ± SD, three independent experiments, asterisks indicate statistical significance by an unpaired two-tailed Student’s *t*-test (*\*p* < 0.05, *\*\*p* < 0.01 and *\*\*\*p* < 0.001). **(C)** The effect of mutation of the IPA element within the *Slp1* promoter on GFP expression in rice sheath cells. Each strain was inoculated and GFP fluorescence was observed at 32 hpi. Scale bar indicates 20 μm.

### *MoHTR1* promoter induces *in planta* expression of a gene specifically expressed during vegetative growth and affects infection-related phenotypes

To determine whether the IPA element of *MoHTR1* can induce the *in planta* expression of other genes, we chose the *MobZIP14* gene encoding a transcription factor involved in conidial morphogenesis and pathogenicity. Notably, *MobZIP14* is highly expressed during vegetative growth but not during infection (Kong et al., 2015), making it an ideal candidate for this expression study. We fused the native promoters of *MobZIP14* and *MoHTR1* to the coding sequence of *MobZIP14* and introduced each construct (*P_MobZIP14_:MobZIP14* and *P_MoHTR1_:MobZIP14*) into *M. oryzae* strain lacking *MobZIP14* **(Fig. 3A and Supplemental Table 1)**. We then compared the *MobZIP14* expression driven by each promoter at the vegetative growth and infection stages using qRT-PCR analysis. During the vegetative growth, the expression of *MobZIP14* in *P_MobZIP14_:MobZIP14* was similar to the wild type, whereas it was reduced by more than 4.6-fold in *P_MoHTR1_:MobZIP14* strain **(Fig. 3B and Supplemental Table 1)**. To observe their expression during infection, we conducted spray inoculation on 4-week-old rice seedlings using wild type, *P_MobZIP14_:MobZIP14* and *P_MoHTR1_:MobZIP14* and collected infected leaves at 48 hpi. In *P_MoHTR1_:MobZIP14*, expression of *MobZIP14* was increased by over 8.7-fold compared with the wild type. However, there was no significant difference in *MobZIP14* expression between *P_MobZIP14_:MobZIP14* and the wild type **(Fig. 3B and Supplemental Table 1)**. In a previous report, *MobZIP14* was shown to be responsible for conidial morphology, conidiation, conidial germination, and appressorium formation (Kong *et al*., 2015). To further investigate whether the *MoHTR1* promoter also affects the roles of *MobZIP14* in fungal development, we assayed conidial morphology, conidiation, conidial germination, and appressorium formation. The *P_MoHTR1_:MobZIP14* strain displayed the phenotypes similar to *ΔMobzip14*, whereas phenotypes of *P_MobZIP14_:MobZIP14* were comparable to WT (**Fig. 3C-F** and **Supplemental Data 2**). These results indicate that the *MoHTR1* promoter, containing the IPA element, can induce *in planta* expression of a gene specifically expressed during vegetative growth and also affect fungal development.

**Figure 3.**
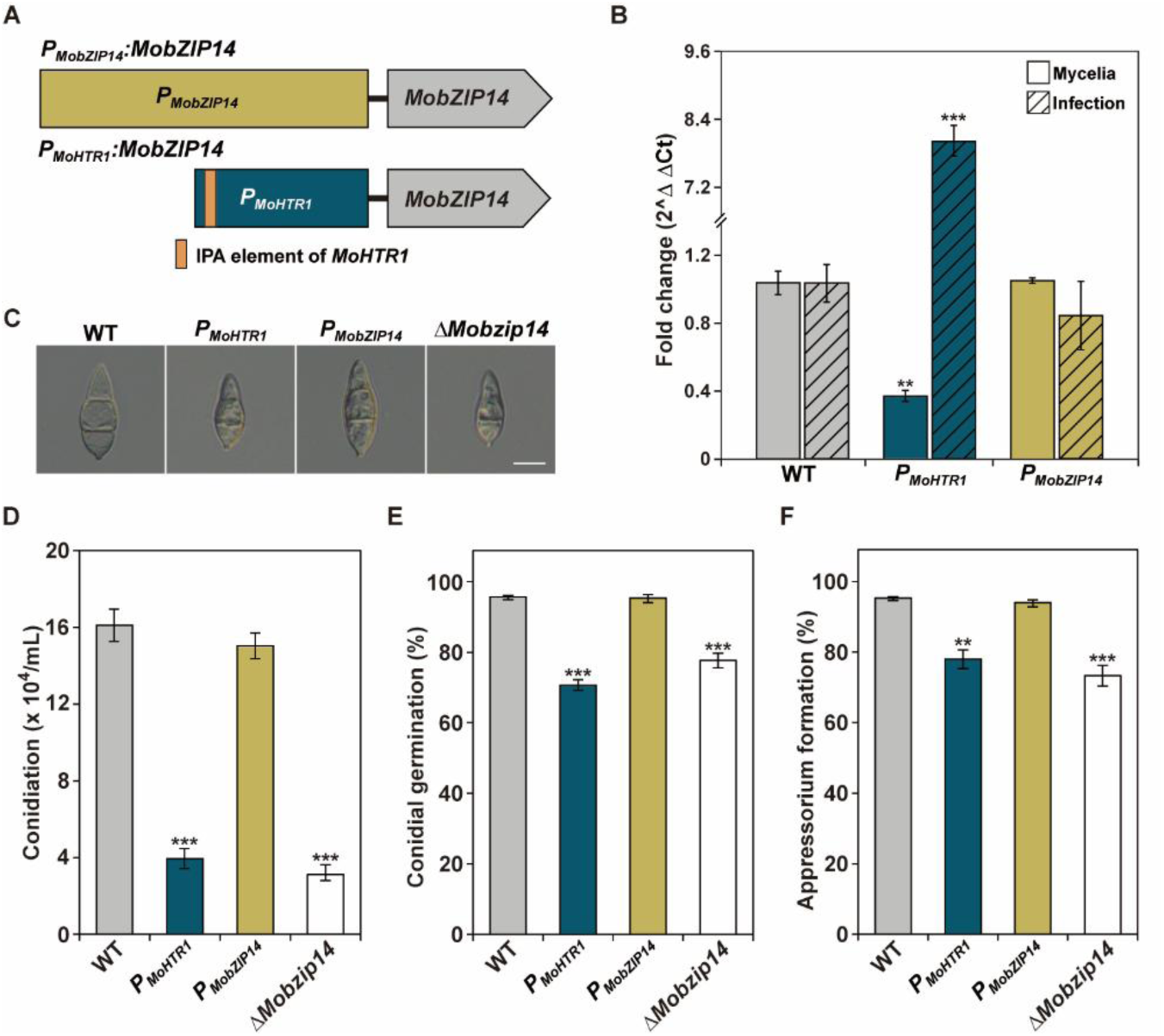
*MoHTR1* promoter induces expression of *MobZIP14* at the infection stage. **(A)** Schematic representation of the promoter switching assay. The *MobZIP14* promoter was swapped with the *MoHTR1* promoter including the IPA element. **(B)** Comparison of transcription profiles of *MobZIP14* according to promoters at the mycelia and infection stages. RNA samples were extracted from mycelia and infected leaves at 48 hpi of WT, *P_MoHTR1_:MobZIP14*, and *P_MobZIP14_:MobZIP14*. Relative expression was calculated using the 2^−ΔΔCt^ method with WT expression normalized to 1.0 for each developmental stage. Mean ± SD, three independent experiments, asterisks indicate statistical significance by an unpaired two-tailed Student’s *t*-test (*\*\*p* < 0.01 and *\*\*\*p* < 0.001). **(C)** Microscopic observation of conidial morphology of WT, *P_MoHTR1_:MobZIP14*, *P_MobZIP14_:MobZIP14*, and *ΔMobzip14*. Conidia were harvested from 7-day-old cultures on V8 agar medium. Scale bar indicates 10 μm. **(D)** Conidia were collected from 7-day-old cultures on V8 agar medium. The number of conidia was measured using a hemocytometer under a microscope. Mean ± SD, three independent experiments, significance was determined by an unpaired two-tailed Student’s t-test (*\*\*\*p* < 0.001). **(E)** Frequency of conidial germination was measured on a hydrophobic surface after 2 h of incubation. Mean ± SD, three independent experiments, significance was determined by an unpaired two-tailed Student’s t-test (*\*\*\*p* < 0.001). **(F)** Frequency of appressorium formation was measured on a hydrophobic surface 8 h of incubation. Mean ± SD, three independent experiments, significance was determined by an unpaired two-tailed Student’s t-test (*\*\*p* < 0.01 and *\*\*\*p* < 0.001).

### The IPA element affects virulence of the rice blast fungus and host immune response

To determine the role of the IPA element of *MoHTR1* in the virulence of *M. oryzae*, we conducted sheath inoculation assays with wild-type, *MoHTR1_pro425_*, *MoHTR1_IPA_mut_*, and *ΔMohtr1* strains. These assays allow for quantitative comparisons of invasive hyphal proliferation inside rice cells. We introduced constructs containing either the normal or IPA-mutated *MoHTR1* promoter fused to the *MoHTR1* coding sequence (*MoHTR1_pro425_* and *MoHTR1_IPA_mut_*) into the *ΔMohtr1* strain. We found that in *MoHTR1_IPA_mut_* and *ΔMohtr1* strains, only 5 % and 7 % of the infection sites showed type Ⅲ growth, respectively, at 48 hpi. In contrast, the wild-type and *MoHTR1_pro425_* exhibited 21 % and 16 % of the infection sites with type Ⅲ growth, respectively **(Fig. 4A)**. *MoHTR1* contributes to fungal virulence by suppressing the expression of immunity-associated genes (Kim et al., 2020a). Given that the *MoHTR1_IPA_mut_* strain exhibited significantly reduced virulence compared to the wild-type, we examined the expression of immunity-associated genes to further investigate the role of the *MoHTR1* IPA element. Several pathogenesis-related genes (*PR1a*, *PR1b*, *PR2*, and *PR10a*) were significantly up-regulated in both the *ΔMohtr1*-infected rice and *MoHTR1_IPA_mut_*-infected rice compared to the infected rice with wild-type **(Fig. 4B)**. These findings indicate that the IPA element contributes to fungal virulence by regulating *MoHTR1* expression, thereby modulating host immune responses.

**Figure 4.**
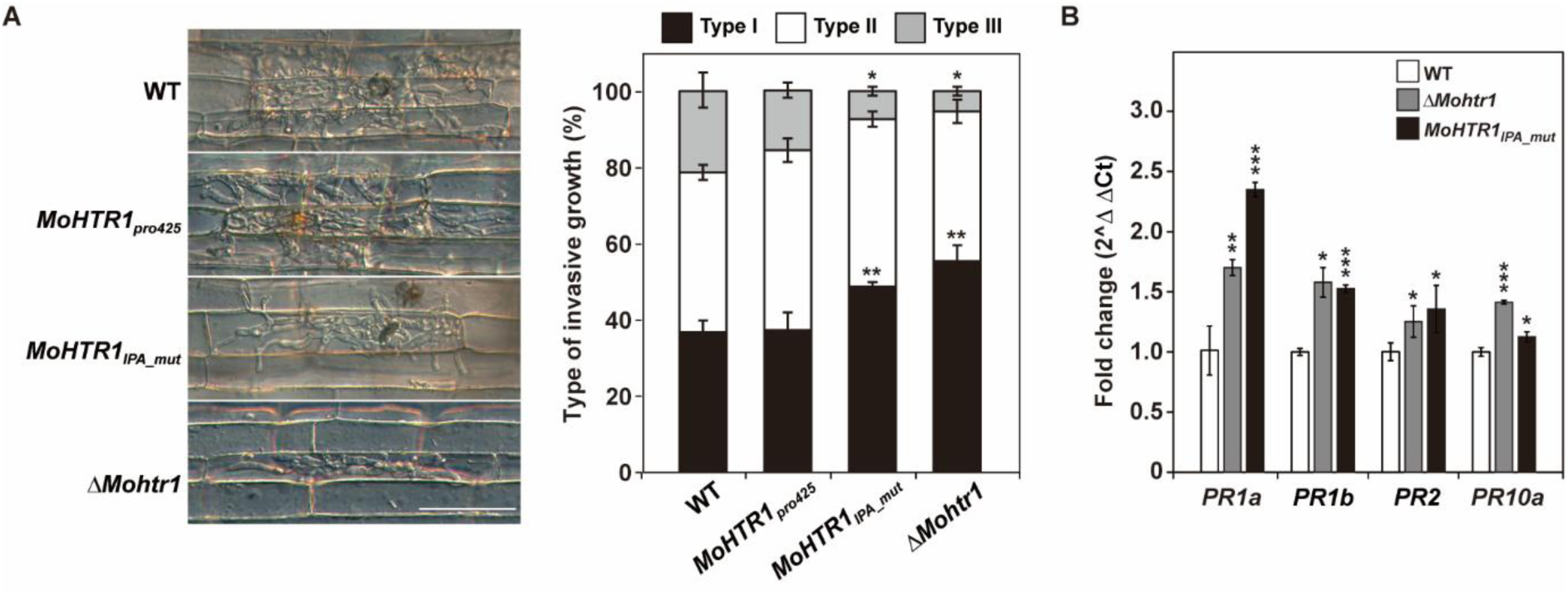
The IPA element of *MoHTR1* is important for the virulence of the rice blast fungus. **(A)** Quantification analysis of the infection severity at 48 hpi. Conidial suspensions (3 × 10^4^ spores/mL) of wild type, *MoHTR1_425pro_*, *MoHTR1_IPA_mut_*, and *ΔMohtr1* were infected into rice sheath cells and observed invasive hyphae under a microscope. Percentages of three different types based on the growth of invasive hyphae were counted at 50 sites of each strain. The criteria for the types are described in Methods. Scale bar indicates 50 μm. Mean ± SD, three independent experiments, asterisks indicate statistical significance by an unpaired two-tailed Student’s *t*-test (*\*p* < 0.05 and *\*\*p* < 0.01). **(B)** Expression levels of pathogenesis-related genes in rice leaves with wild type, *ΔMohtr1*, and *MoHTR1_IPA_mut_* strains. Infected rice leaves were collected at 48 hpi after spray inoculation. The abundance of transcripts of pathogenesis-related genes was quantified using qRT-PCR. Relative expression was calculated using the 2^−ΔΔCt^ method with wild type-infected samples as the reference, and *Actin* was used for normalization. Mean ± SD, three independent experiments, asterisks indicate statistical significance by an unpaired two-tailed Student’s *t*-test (*\*p* < 0.05, *\*\*p* < 0.01 and *\*\*\*p* < 0.001).

### Eight transcription factors interact *in vitro* with the IPA element of *MoHTR1*

To identify transcription factors that regulate *MoHTR1* expression by binding to its IPA element, we employed two strategies: profiling based on gene expression patterns and performing a pull-down assay. First, from the 495 predicted transcription factors in *M. oryzae*, we selected the candidate activators with the same expression pattern as *MoHTR1* and candidate suppressors with opposing expression pattern using RNA-seq data (Jeon *et al*., 2020; Park et al., 2013). Four activator and four suppressor transcription factors were chosen for subsequent interaction assay **(Supplemental Data 3)**. Second, we performed a pull-down assay using two biotinylated oligonucleotide probes (IPA probe and IPA mutant probe). After incubating each probe with total protein extracted from *M. oryzae* mycelia, we identified 45 candidate transcription factors that bound to the IPA probe, but not to the mutant probe via liquid chromatography-mass spectrometry (LC-MS) **(Supplemental Table 2)**. We then selected 14 previously characterized transcription factors that were also detected in at least two replicates **(Supplemental Data 3)**.

To further validate the interactions between transcription factors with the IPA element, we conducted a yeast-one hybrid (Y1H) assay. We cloned a 40-bp *MoHTR1* promoter containing the IPA element (IPA) and a transversion-mutated IPA region (IPA-mutant), along with five copies of the IPA element (5 × IPA) into bait vector. We also cloned the coding sequences of 22 transcription factors (eight identified through RNA-seq profiling and 14 identified from a pull-down assay) into the prey vector **(Fig. 5A)**. Among these 22 transcription factors **(Supplemental Data 3)**, eight transcription factors bound to both the IPA and 5 × IPA baits, but not to the IPA mutant bait **(Fig. 5B and Supplemental Fig. 5).** These include four previously characterized transcription factors: RGS1 (winged-helix binding domain), RPP3 (Zn2Cys6), MoHOX3 (Homeobox), and MobZIP15(bZIP) (Cao *et al*., 2022; Kim et al., 2009; Kong *et al*., 2015; Osés-Ruiz et al., 2021). These results suggest that eight transcription factors of *M. oryzae* may regulate *MoHTR1* expression by directly binding to the IPA element.

**Figure 5.**
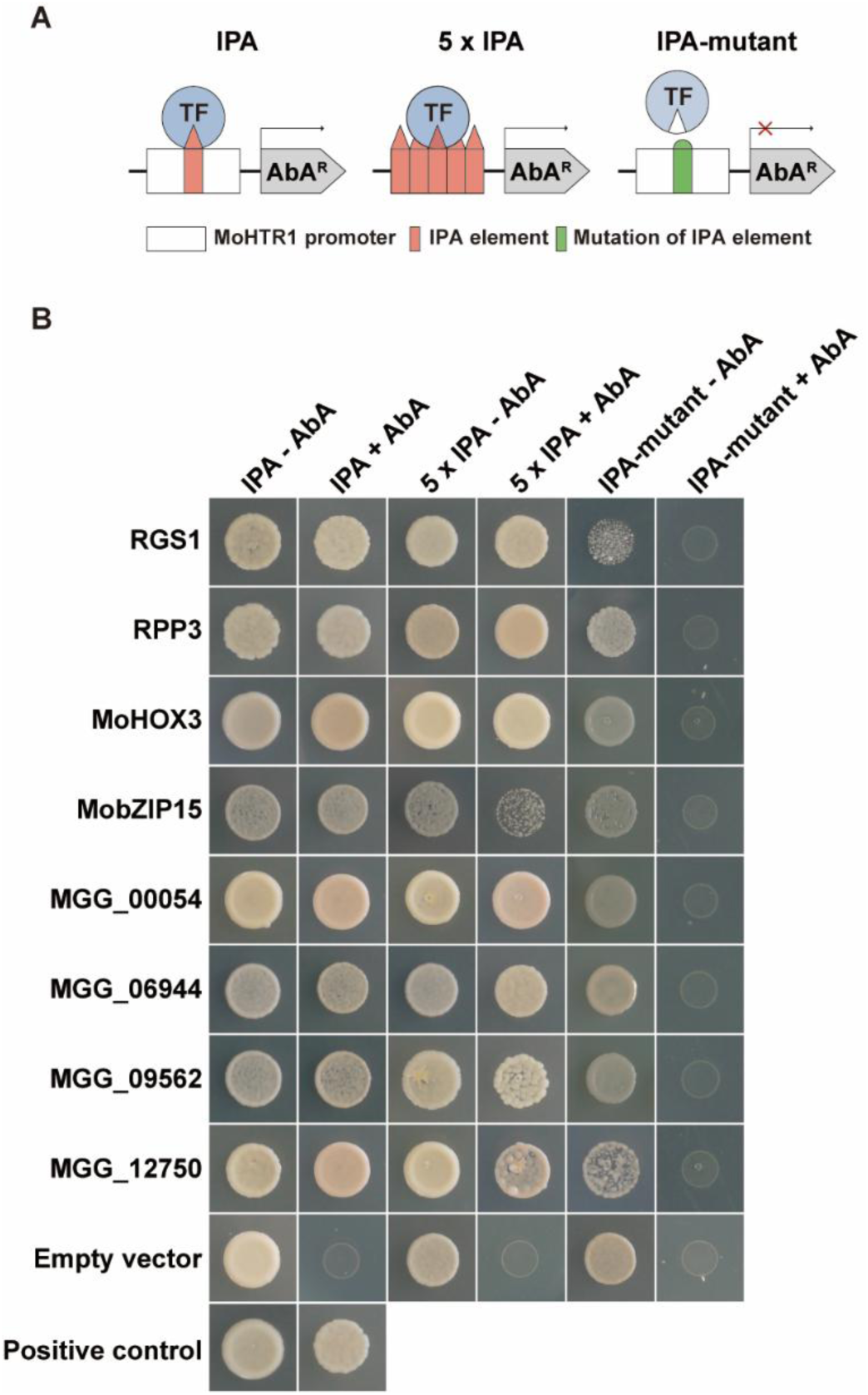
Transcription factor candidates binding to the IPA element of *MoHTR1* were selected using Y1H assay. **(A)** Schematic diagrams of baits used in Y1H assays. Three different fragments (IPA: -442 to -402 region from the start codon of *MoHTR1*, 5 × IPA: five times repeat sequence of the 8-bp IPA element, and IPA mutant: transversion mutation of 8-bp of the IPA element sequence among 40-bp of promoter regions) were inserted into the pAbAi vector. Selected transcription factor candidates were cloned into the prey vector, pGADT7. **(B)** Analysis of TF binding to the IPA element by Y1H assay. pGADT7Rec-p53/p53-AbAi was used as the positive control. Bait-only cells carrying pGADT7 (empty vector, EV) were plated on SD/−Leu medium containing AbA (aureobasidin A) to assess basal auto-activation. No growth was observed at the working AbA concentration. To test interactions, the bait and prey constructs were co-transformed into the Y1H Gold yeast strain and grown on SD/−Leu medium containing AbA. AbA was used as a screening marker.

### RGS1 acts as a transcriptional regulator of genes containing the IPA element

Before conducting an in-depth analysis of the interactions between the *MoHTR1* IPA element and transcription factors, we used AlphaFold analysis to identify those with higher binding potential. Among eight transcription factors identified through the Y1H assay, four previously characterized ones showed strong interaction potentials **(Supplemental Fig. 6A)**. To further confirm the interaction, we focused on RGS1 which was identified as a regulator of effector gene expression during conidiation. Consistent with a previous report, we observed that the *Δrgs1* mutant exhibited defects in mycelial growth, conidiation, conidial germination, appressorium formation, and pathogenicity **(Supplemental Fig. 7)**. Next, we carried out an electrophoretic mobility shift assays (EMSA) using recombinant RGS1 protein with 20-bp biotin-labeled oligonucleotides containing the IPA element sequence. The EMSA results showed a shifted band when the purified 6 × His RGS1 fusion protein was incubated with the biotinylated probe containing the IPA element. In contrast, no mobility shift was detected when RGS1 was incubated with a high concentration of non-biotinylated competitive probe **(Fig. 6A)**. These results further support that RGS1 directly binds to the IPA element of *MoHTR1*, consistent with previous data from the Y1H assay. To address how RGS1 binds to the IPA element, we performed further computational analyses. Domain annotation with InterProScan identified a winged-helix-like DNA-binding domain (DBD) spanning from amino acids 210 to 498. AlphaFold analysis showed that this DBD interacts with the IPA element with higher confidence (average PAE = 17.9) than the full-length RGS1 (average PAE = 25.0). These results suggest that IPA element is directly recognized by the RGS1 DNA-binding domain **(Supplemental Fig. 6B)**. To examine how RGS1 regulates *MoHTR1* expression, we first analyzed *RGS1* expression levels during infection stages, comparing them to the mycelial growth stage. *RGS1* expression was highly induced during infection stages at 24, 48, and 72 hpi, consistent with high expression of *MoHTR1* at these stages **(Fig. 6B)**. We also examined *MoHTR1* expression in the wild-type and *Δrgs1* strains during the mycelial growth and infection stages. RNA samples were collected from the mycelia and *M. oryzae*-infected leaves of each strain at 24, 48, and 72 hpi. *MoHTR1* expression decreased significantly in all infected samples of the *Δrgs1* strain compared to the wild type, with a 7.69-fold decrease at 48 hpi, when *MoHTR1* expression peaked **(Supplemental Fig. 1A and Fig. 6C).** In the *Δrgs1* strain, however, *MoHTR1* was up-regulated by 2.41-fold compared to the wild type in the mycelial growth stage **(Fig. 6C)**. These results indicate that RGS1 suppresses *MoHTR1* expression during mycelial growth but activates it during the infection stages. We found a similar regulation for *Slp1*, with RGS1 suppressing its expression in the mycelial stage and activating it during the infection stages **(Fig. 6D)**. Given the shared presence of the IPA element in the promoters of *MoHTR1* and *Slp1*, our findings suggest that RGS1 directly binds to the IPA element to coordinate the expression of both genes during infection.

**Figure 6.**
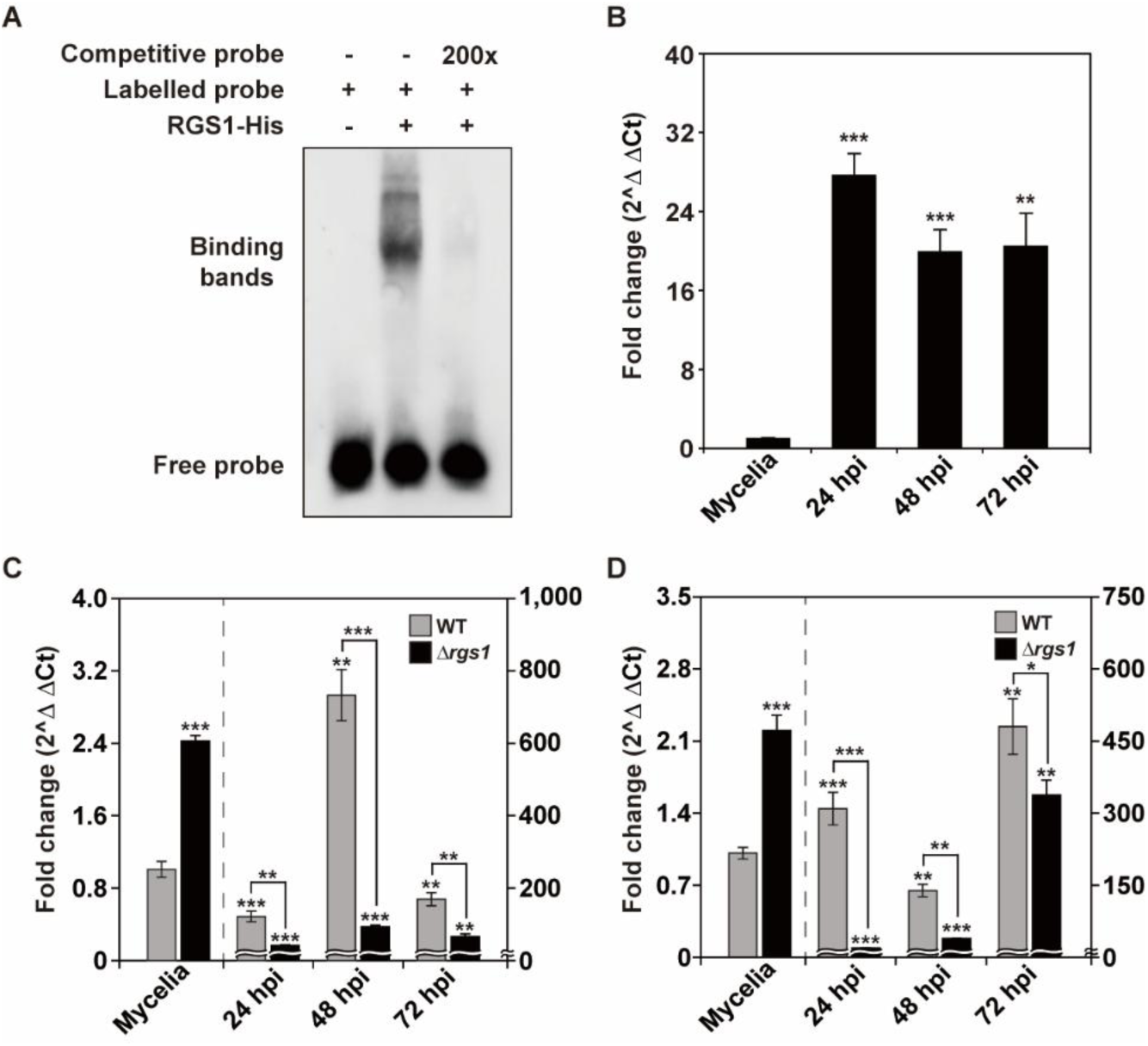
RGS1 directly binds to the IPA element of *MoHTR1* and regulates expression of *MoHTR1*. **(A)** RGS1 binds directly to the IPA element of *MoHTR1* in electrophoretic mobility shift assay (EMSA). A biotin-labeled probe containing the IPA element was incubated with RGS1-His protein. The 200-fold molar excess of unlabeled probes was used as competitors. **(B)** Expression profile of *RGS1* during the mycelial growth and infection stages. Relative expression was calculated using the 2^−ΔΔCt^ method with mycelial samples as the reference. Mean ± SD, three independent experiments, asterisks indicate statistical significance by an unpaired two-tailed Student’s *t*-test (*\*\*p* < 0.01 and *\*\*\*p* < 0.001). **(C)** Comparison of expression patterns of *MoHTR1* at the various infection stages. RNA samples were extracted from mycelia and infected leaves at 24, 48, and 72 hpi of WT and *ΔMorgs1* respectively. Transcript levels were normalized using *β-tubulin* and all values were expressed relative to WT mycelium as the reference condition using the 2^−ΔΔCt^ method. Mean ± SD, three independent experiments, asterisks indicate statistical significance by an unpaired two-tailed Student’s *t*-test (*\*\*p* < 0.01 and *\*\*\*p* < 0.001). **(D)** Comparison of expression patterns of *Slp1* at mycelia and infection stages. RNA samples were harvested from mycelia, and infected leaves at 24, 48, and 72 hpi of WT and *ΔMorgs1*, respectively. Transcript levels were normalized using β-tubulin gene and all values were expressed relative to WT mycelium as the reference condition using the 2^−ΔΔCt^ method. Mean ± SD, three independent experiments, asterisks indicate statistical significance by an unpaired two-tailed Student’s *t*-test (*\*p* < 0.05, *\*\*p* < 0.01 and *\*\*\*p* < 0.001).

## DISCUSSION

Effectors are secreted into host plant cells to suppress host immunity and facilitate pathogen infection. Their expression is tightly regulated during various developmental and infection stages to ensure successful colonization. While effector functions have been extensively studied, the transcriptional regulatory mechanisms governing effector gene expression remain poorly understood. In this study, we identified the core *cis*-regulatory IPA (*in planta* active) element responsible for the *in planta*-specific expression of *MoHTR1.* This expression is induced by an unidentified, viable plant-derived signal common to both host and non-host plants. Furthermore, we demonstrated that the transcription factor RGS1 directly binds to the IPA element to regulate *MoHTR1* expression **(Fig. 7)**.

**Figure 7.**
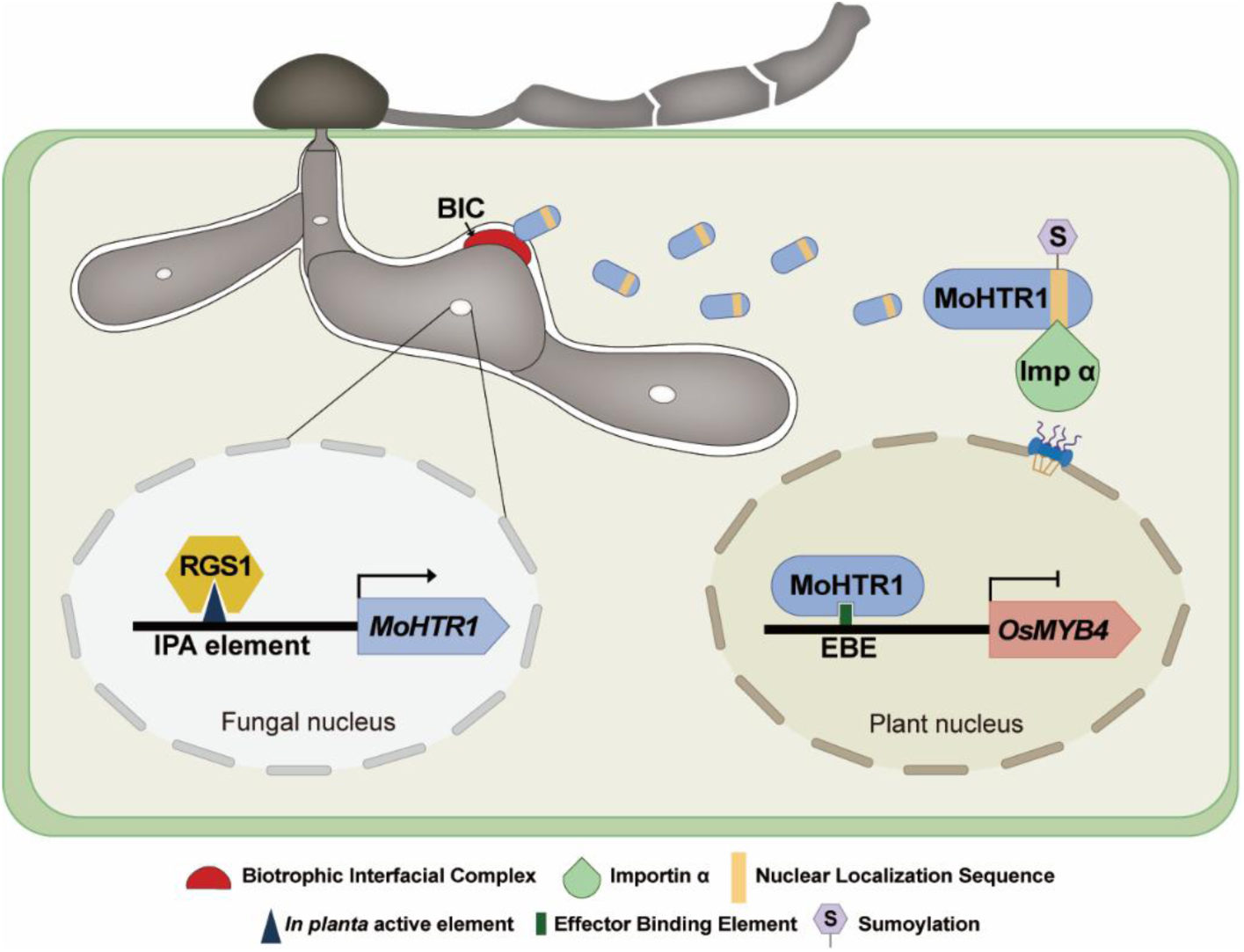
The proposed model for the transcriptional regulatory mechanism of effector genes. During *M. oryzae* infection, RGS1 acts as an activator transcription factor by directly binds to the IPA element in the *MoHTR1* promoter. Consequently, MoHTR1 is secreted into the host cell via BIC. For translocation of MoHTR1 to host nuclei, the core nucleus localization sequence (NLS) of MoHTR1 interacts with importin α protein. SUMOylation is crucial for nuclear localization of MoHTR1. In plant nucleus, MoHTR1 modulate host immunity through suppressing the expression of *OsMYB4*, a host immunity-associated gene.

While several studies have identified *cis*-elements in the promoters of effector genes associated with *in planta*-specific expression in bacteria and Oomycetes, research on *cis*-elements in plant pathogenic fungi is very limited. The plant-inducible promoter (PIP) box, found in the promoters of bacterial type III effector genes, is conserved across species such as *Xanthomonas campestris* pv. *versicatoria*, *Burkholderia sensu lato*, and *Ralstonia solanacearum* (Cunnac *et al*., 2004; Koebnik et al., 2006; Wallner et al., 2021). Similarly, genes containing the AM-20 *cis*-element are highly expressed during infection and are conserved in the Oomycete pathogens including *Phytophthora infestans*, *P. sojae*, and *P. ramorum* (Roy et al., 2013; Seidl et al., 2012). In this study, we identified an 8-bp *cis*-element (TATTTCGT), which is crucial for the *in planta*-specific expression of *MoHTR1*. This was demonstrated through serial deletion of the promoter region of *MoHTR1* and transversion mutation of the IPA element, combined with live cell imaging experiments. When the PIP box-like sequence within the IPA element was mutated, promoter activity was significantly reduced. Additionally, mutation of remaining region of the IPA element-excluding the PIP box-like sequence-completely abolished promoter activity. In the case of the PIP box (TTCG-N_16_-TTCG), both conserved TTCG repeats and distance between TTCG sequence are essential for promoter activity (Cunnac *et al*., 2004; Tsuge et al., 2005). These data strongly suggest that the second TTCG sequence, separated by 154 nucleotides, is unlikely to contribute to its promoter function in the *MoHTR1* promoter. These findings indicate that the IPA element functions as a core *cis*-element that is distinct from previously reported *cis*-elements in bacterial and Oomycete pathogens. When we searched this IPA element sequence on the genome of *M. oryzae*, 234 genes which are highly expressed during infection were identified. Among these, the IPA element sequence was present in one well-characterized apoplastic effector gene, *Slp1*, and other four effector candidate genes. Furthermore, qRT-PCR analysis confirmed that these effectors are all highly expressed during infection. These data suggest that the IPA element sequence is responsible for *in planta*-specific expression of not only *MoHTR1* but also other effector genes. Recently, a *cis*-element (TTATGCAAGCTT) associated with the biotrophic stage-specific expression of *PWL2*, a cytoplasmic effector gene of the rice blast fungus, was reported (Zhu et al., 2021). However, the regulatory mechanism for *PWL2* during the biotrophic stage and whether other genes containing this *cis*-element are controlled by this element remains unclear. The functionality of the IPA element for *in planta*-specific expression was further validated through two complementary experiments. First, transversion mutation of the IPA element in the *Slp1* promoter completely abolished its expression during infection. Second, when the promoter of *MobZIP14*, which encodes a bZIP transcription factor and is specifically expressed during vegetative growth, was replaced with the promoter of *MoHTR1* containing the IPA element, expression of *MobZIP14* was strongly induced during infection. These data strongly support that the IPA element is responsible for *in planta*-specific expression of *MoHTR1* and other effector genes.

Many transcription factor have been characterized in plant pathogenic fungi, but studies focusing on the regulatory mechanism of effector genes remain limited. For instance, PnCon7 in *Parastagonospora nodorum* regulates *Tox3* expression, but its precise *cis*-element remains unidentified (Lin et al., 2018). Similarly, PsMyb37 in *Phytophthora sojae* binds to the AM-20 *cis*-element (TACATGTA) in the promoters of *PsAvr3c* and *PsAvr1k*, regulating their expression during infection (Qian et al., 2024). Our study extends these findings and provides new insights into the regulatory mechanisms of the *M. oryzae* transcription factor RGS1, specifically targeting the regulation of effector genes containing the IPA element. The role of RGS1 as a positive transcriptional regulator of *MoHTR1* expression was confirmed using a deletion mutant of RGS1. In the *Δrgs1* strain, transcriptional activation of *MoHTR1* was significantly compromised in invasive hyphae during rice infection. We further demonstrated the direct binding of RGS1 to the IPA element using yeast one-hybrid and electrophoretic mobility shift assays. Consistently, mutation of the IPA element abolished *MoHTR1* expression in invasive hyphae, even in the presence of RGS1, underscoring the critical role of the IPA element and RGS1 in effector gene activation. Notably, our data reveal a dual regulatory function of RGS1; it acts as a repressor of *MoHTR1* and *Slp1* during pre-penetration stages, as evidenced by transcriptional activation in vegetative hyphae upon RGS1 loss, whereas it functions as an activator during infection stages. This dual regulatory pattern was consistently observed in the regulation of both *MoHTR1* and another IPA element-containing gene, *Slp1*, highlighting RGS1’s critical role in temporal coordination of effector expression. Previously, Tang *et al*. (2023) reported that RGS1 represses over 60 effector genes during the pre-penetration stages of development before plant infection. However, *MoHTR1* and *Slp1* were not included among these genes. One possible explanation is that their RNA-seq analysis, conducted without knowledge of the IPA element, focused specifically on the conidiation stage, during which *MoHTR1* and *Slp1* are rarely expressed. While their work primarily highlighted RGS1’s repressive role during pre-penetration, our findings reveal an additional activation function during the post-penetration stage for effector genes regulated by the IPA element. Importantly, our study reveals a distinct regulatory mechanism from previously characterized RGS1-regulated effectors. Our analysis showed that seven previously reported RGS1-repressed effector genes, including *Mep2*, do not contain the IPA element in their promoter regions, even when extending the search up to 2 kb upstream of the TSS. This observation is consistent with a prior study suggesting that RGS1 regulates these genes through indirect mechanisms. In contrast, our results support a distinct mechanism in which RGS1 directly binds the IPA element to activate a subset of effectors during infection, including *MoHTR1* and *Slp1*. The underlying mechanisms of this stage-specific switch between repression and activation remain unclear, such as whether RGS1 directly binds to the IPA element when acting as a repressor, adding complexity to its regulatory functions. Nevertheless, the regulatory system involving the IPA element of target effector genes and RGS1 offers a valuable model for further exploration. It is also notable that the MoHTR1 and Slp1 proteins play distinct roles in modulating host immunity, each acting at a different subcellular location after their secretion from invasive hyphae—Slp1 within the EIHM compartment and MoHTR1 in the rice nucleus. Taken together, we propose a model in which RGS1 dynamically regulates effector gene expression through stage-specific interactions with the IPA element, ensuring precise temporal coordination critical for fungal virulence.

In addition to RGS1, three other TFs (RPP3, MoHOX3, and MobZIP15) also showed high affinity for the IPA element. A combinatorial regulatory model in which multiple TFs compete or cooperate at a shared *cis*-element to regulate gene expression has been reported in plants. For instance, HY5 and PIF can compete for G-box motifs (Toledo-Ortiz et al., 2014), whereas MYC2/3/4 can act additively at G-box motifs (Zhang et al., 2020). In this context, we cannot completely exclude the possibility that, in addition to Rgs1, other IPA-binding TFs may compete or cooperate at the IPA element to regulate gene expression.

Expression of *MoHTR1* was induced only when inoculated into living cells of both host (rice and barley) and non-host (onion) plants, but it failed to be induced in dead plant cells. These results demonstrate that *MoHTR1* is specifically expressed during infection stages and suggest that this *in planta*-specific expression is induced by a vital signal(s) conserved across plant species. Upon pathogen infection, multiple signaling pathways are rapidly activated in living plant cells (Sewelam et al., 2016). In addition, damage-associated molecular patterns (DAMPs), such as wax and cutin monomers, are synthesized and accumulated in viable epidermal cells during infection (Yeats and Rose, 2013; Ziv et al., 2018). However, our results demonstrate that none of these responses, including ROS accumulation, Ca2+ flux, MAP kinase cascade, and SA, and JA signalings as well as cutin monomer biosynthesis, are responsible for inducing *in planta*-specific expression of *MoHTR1*. These findings suggest that a vitality-dependent plant signal(s), distinct from known immune signaling pathways and cutin-derived molecules, are essential for triggering *MoHTR1* expression *in planta*.

In summary, we identified the core IPA element sequence responsible for *in-planta* specific expression of *MoHTR1* and other effector genes in the rice blast fungus. Additionally, we also confirmed that RGS1, a transcription factor containing a winged-helix binding domain, acts as a regulator of *MoHTR1* expression by directly binding on the IPA element and contributes to pathogenicity of *M. oryzae*. Our study provides the first comprehensive regulatory mechanism of effector gene expression, including those secreted into host cells via BIC, as well as those secreted between invasive hyphae and the EIHM in the rice blast fungus. Building on these findings, future research should aim to identify additional transcription factors and explore host-derived signaling molecules that interact with this pathway. This would provide deeper insights into the coordination of *in-planta* specific expression across diverse effector genes, offering potential strategies for management of fungal diseases.

## METHODS

### Plasmid construct for promoter activity analysis

To perform promoter truncation analysis, we amplified the promoter region (900-bp upstream of *MoHTR1* excluding the low complexity poly (T) stretch from -969 to -917) from the genomic DNA (gDNA) of *Magnaporthe oryzae* wild type strain, KJ201. And then, we truncated promoter region to -900, -750, -600, -450, -425, -400, -350, and -300 bp upstream of *MoHTR1*. To identify the target IPA element region, we conducted transversion substitution of 12-bp in three candidates of the IPA motif (IPA motif1: -425 to -415 region, IPA motif2: -419 to -409 region, IPA motif3: -412 to -401 region from the start codon of *MoHTR1*). To verify the IPA element of *MoHTR1* that regulates the expression of *Slp1*, we amplified the native promoter of *Slp1* (*Slp1_pro815_*) from gDNA of KJ201 and generated the transversion mutated promoter of *Slp1* (*Slp1_IPA_mut_*). Each upstream region and transversion mutated IPA element candidates were tagged with sGFP and cloned into pCB1004 vector. After then, Each plasmids was transformed into KJ201 protoplast using PEG-mediated transformation (Liu and Friesen, 2012). All transformants for promoter activity analysis were selected on TB3 agar media (0.3% yeast extract, 0.3% casamino acid, 1% glucose, 20% sucrose, and 0.8% agar) containing 200 µg/mL of Hygromycin B and were confirmed by mycelial PCR using designed primers **(Supplemental Table 3)**.

### Microscopic observation and image analysis

For a precise comparison of fluorescence signals, promoter activities were assessed in five strains (*MoHTR1_pro425_*, *MoHTR1_pro400_*, *MoHTR1_IPA1_mut_*, *MoHTR1_IPA2_mut_*, and *MoHTR1_IPA3_mut_*), each carrying a different promoter construct fused to sGFP. To examine whether the IPA element functions differently from the bacterial PIP box, two additional mutant constructs were generated: *MoHTR1_IPA1_mut1_*, in which the PIP box-like region within the IPA element was transversion-mutated, and *MoHTR1_IPA1_mut2_*, in which the remaining region excluding the PIP box-like sequence was mutated. Each construct was introduced into KJ201 protoplast.

For subcellular localization under promoters with mutations in the IPA element, the native promoter and two IPA element-mutated promoters were inserted into the pFPL-Rh vector, respectively (Gong et al., 2015). These constructs were co-transformed with PWL2pro:PWL2:eGFP:NLS to label BICs and rice nuclei into KJ201 protoplasts.

For promoter activity assays, we did sheath inoculation with conidia suspension (3 × 10^4^/mL in sterile water) of each transformants into 6-week-old rice seedlings (*Orayza sativa* cv. Nakdongbyeo) and observed using a fluorescence microscope at 32 hpi. To determine whether host range and plant viability is required for the *in planta*-specific expression of *MoHTR1*, we inoculated a conidial suspension of the transformant co-expressing EF1αpro:mRFP and MoHTR1pro:sGFP into various plant tissues. These included live sheath cells of 6-week-old rice seedlings, epidermis cells of 1-week-old barley seedlings and onion bulbs, as well as heat-killed seedlings that had been incubated at 70 ℃ for 25 min. To further elucidate the involvement of pathogen infection-derived plant signaling pathways in *MoHTR1* expression, we inoculated a conidial suspension with specific inhibitors. Each inhibitor was applied at a range of concentrations, from levels that allowed invasive hyphae (IH) development comparable to the DMSO-treated control, to levels that completely inhibited the formation of germ tube or appressorium. These included diphenyleneiodonium chloride (DPI; 0.4-4 μM) to block ROS accumulation, BAPTA-AM (20-100 Μm) to inhibit Ca^2+^ flux, U0126 (5-50 μM) to suppress MAP kinase cascades, 2-aminoindan-2-phosphonic acid (AIP; 30-200 μM) to interfere with salicylic acid (SA) signaling, and phenidone (50-300 μM) to inhibit jasmonic acid (JA) signaling. In addition, to assess the role of cutin monomers in *MoHTR1* expression, we treated 100 μM of each cutin monomers, including 1,16-Hexadecanediol (1,16-HDC), 12-Hydroxystearic acid (12-HSA), and 16-Hydroxyhexadecanoic acid (16-HDA), with a conidial suspension of the transformant co-expressing EF1αpro:mRFP and MoHTR1pro:sGFP. Microscopic observation was performed using Leica DM6 B microscope, and taken images using Leica DMC6200 camera (Leica Microsystems). Excitation was 480/40 nm for GFP and images were processed using LAS X software. Promoter activity was quantified by measuring GFP intensity at 20 infection sites per strain using ImageJ software.

### RNA extraction and qRT-PCR

To investigate whether the *MoHTR1* promoter affects the expression patterns of *MobZIP14*, we performed the promoter switching experiment. We designed *MobZIP14* constructs expressed under the native *MobZIP14* promoter and *MoHTR1* promoter (*P_MobZIP14_:MobZIP14* and *P_MoHTR1_:MobZIP14*) and introduced these into *ΔMobZIP14* protoplasts. The *ΔMobZIP14* strain was generated in a previous study (Kong et al., 2015). To confirm the expression of *MobZIP14* during the infection stage, we conducted spray inoculation on 4-week-old rice seedlings with conidial suspensions (1 × 10^5^/mL) of the wild type, *P_MobZIP14_:MobZIP14*, and *P_MoHTR1_:MobZIP14*. The infected rice leaves were harvested at 48 hpi.

To examine whether the IPA element regulates the expression of host immunity-associated genes during infection, we generated the *MoHTR1_IPA_mut_* strain by introducing a construct containing a mutated IPA element into the *ΔMohtr1* protoplast. To compare the expression levels of pathogenesis-related genes, conidial suspensions (5 × 10^5^/mL) of the WT, *ΔMohtr1*, and *MoHTR1_IPA_mut_* strains were spray-inoculated onto 4-week-old rice seedlings, and infected leaves were collected at 48 hpi.

To identify how RGS1 regulates the expression of *MoHTR1* at mycelial growth and infection stages of *M. oryzae*, we collected *M. oryzae* samples and extracted total RNA from four stages (mycelial growth, 24 hpi, 48 hpi, and 72 hpi). Mycelia were cultured in CM broth medium (6 % yeast extract, 6 % casamino acids, and 10 % sucrose) at 25 °C for 3 days. To collect the infection stage samples, 10 mL conidial suspension (5 × 10^5^/mL) containing Tween 20 (250 ppm) was sprayed on 4-week-old rice seedling, and infected leaves were harvested at 24, 48, and 72 hpi.

Total RNA was extracted using the Easy-spin total RNA extraction kit (iNtRON Biotechnology) according to the manufacturer’s instructions. cDNA was synthesized using ImProm-II Reverse Transcription System (Promega) from 5 μg of total RNA. For qRT-PCR experiment, we used a Rotor-Gene Q real-time PCR cycler (Qiagen) and TOPreal™ qPCR 2X PreMIX (Enzynomics). Each reaction tube contained 5 μl of TOPreal™ qPCR 2X PreMIX, 2 μL of cDNA (12.5 ng/μL), and 1.5 μL of each forward and reverse primers (100 nM for each primer). The primers for qRT-PCR are listed in the **Supplemental Table 3**. All qRT-PCR amplifications followed the thermal cycling conditions: 3 min at 95 °C; followed by 40 cycles of 95 °C for 15 sec, 60 °C for 30 sec, and 72 °C for 30 sec.

### Pathogenicity test

To confirm whether the IPA element of *MoHTR1* affects *in planta*-specific expression as well as virulence, we performed a sheath inoculation assay. We generated *MoHTR1* constructs, expressed under *MoHTR1* promoter with an intact IPA element (*MoHTR1_pro425_*) and with a transversion mutated IPA element (*MoHTR1_IPA_mut_*), and introduced each construct into Δ*Mohtr1* protoplast. The Δ*Mohtr1* strain was generated in a previous study (Kim et al., 2020a). In addition, to perform pathogenicity test of wild type, Δ*rgs1*, and *rgs1c* strains, we spray-inoculated conidial suspensions (5 × 10^4^/mL) of each strain on 4-week-old rice seedlings. Inoculated rice plants were incubated at 25 °C for 24 h at a relative humidity of 100 % in the dark, and then incubated at 28 °C in a growth chamber with 16/8 h light/dark cycle. Infected rice leaves were collected at 6 dpi. For sheath inoculation assay, conidial suspensions (3 × 10^4^/mL) of each strain were inoculated on sheath of 6-week-old rice seedlings. We observed the invasive hyphal (IH) growth of each strain at 50 infection sites at 48 hpi. We categorized the IH growth into three types. Type I for primary IH in the penetrated cell, type II for IH growing to adjacent cells, and type III for IH growing beyond the adjacent cells.

### Pull-down assay

A pull-down assay was performed as previous described (Wu, 2006). Total proteins were extracted from frozen mycelial powder using ice-cold extraction buffer [10 % glycerol, 25 mM Tris (pH 7.5), 1 mM EDTA, 150 mM NaCl, 2 % w/v PVPP, 10 mM dithiothreitol, 1× protease inhibitor cocktail, and 0.1 % Tween 20]. After centrifugation at 3200 g for 30 min at 4 °C, the supernatant was filtered using a syringe filter (0.45 μm). Protein concentration was measured using the Bradford assay. Both 100-bp of sense and antisense biotinylated oligonucleotide (1μg/μL) containing IPA element were incubated at 100 °C for 1 h, and cool down at 25 °C for 1 h. The biotinylated double-stranded oligonucleotides are listed in **Supplemental Table 3**. Extracted proteins were mixed with biotin-labeled probe and streptavidin agarose beads, then incubated for 2.5 h at 25 °C with gentle speed (25 rpm). After incubation, we washed the protein–DNA–streptavidin–agarose complex three times with PBS buffer containing protease inhibitors. Then resuspended in 50 μL of 2× Laemmli sample buffer and incubated at 95 °C for 5 min. After centrifuging at 7000 g for 30 sec, the supernatant was collected and loaded on 12 % Tris-glycine gel and was separated by SDS-PAGE gel electrophoresis. LC-Mass spectrometry was performed at National Instrumentation Center for Environmental Management at Seoul National University (NICEM).

### Yeast-one hybrid

Yeast-one-hybrid assays were performed using the Matchmaker Gold Yeast One-Hybrid Library Screening System (Clontech) according to the manufacturer’s instructions. We generated three kinds of 40-bp of baits (IPA: -442 to -402 region from the start codon of *MoHTR1*, 5 × IPA: five times repeat sequence of 8-bp IPA element, and IPA mutant: transversion mutation of 8-bp of IPA element sequence among 40-bp of promoter regions) and cloned them into the pAbAi vector, respectively. pAbAi:IPA, pAbAi: 5 × IPA, and pAbAi:IPA mutant were linearized and transformed into the yeast strain, Y1H gold via LiAc-mediated transformation. The full-length coding sequence (CDS) of TF candidates were cloned into the pGADT7 vector (prey) with the GAL4 activation domain. All prey plasmids were transformed into the yeast strains carrying each bait-reporter, respectively. The interaction between TF candidates and baits, including IPA, 5 × IPA, and IPA mutant was confirmed using the selection media (SD/-Leu agar media containing 100 ng/mL Aureobasidin A (AbA)).

### Electrophoretic mobility shift assay (EMSA)

To confirm the binding ability of RGS1 to the IPA element of *MoHTR1*, His6 tagged RGS1 was expressed using the pET-28a vector (Novagen) in *Escherichia coli strain* BL21 by adding 0.1 mM IPTG. We purified His-RGS1 using Ni sepharose beads and eluted it using imidazole. After purification, the elution buffer was exchanged with PBS using Amicon Ultra 0.5 centrifugal filters (Merck KGaA). Biotin-labeled and unlabeled 20-bp of DNA probes were synthesized by Bionics Company (Bionics). EMSA binding reactions were performed using the LightShift™ Chemiluminescent EMSA Kit (Thermo Scientific) following the manufacturer’s protocol. Purified His-RGS1 was incubated with biotin labeled or unlabeled probes in binding reaction mixtures for 20 min at 25 °C. After incubation, the reaction mixtures were loaded onto 5 % polyacrylamide gels for 1 h at 100 V, and then transferred to a positively charged nylon membrane (Thermo Scientific). The band shifts were detected by a Chemi-Doc imaging system (Bio-rad).

### AlphaFold analysis

Predictions for protein-DNA interactions were performed with AlphaFold 3 (Abramson et al., 2024). Model quality was evaluated using pLDDT (predicted local distance difference test) and PAE (predicted aligned error) (Elfmann and Stülke, 2023). For protein-DNA complexes, we calculated the mean inter-chain PAE over residues at the predicted protein–DNA interface. Lower PAE values were interpreted as indicating higher confidence in the predicted interaction.

### Target gene deletion and complementation

For deletion of *RGS1*, 1.5-kb upstream and downstream sequences were amplified using designed primers **(Supplemental Table 3)** from the gDNA of KJ201, and then fused to amplified HPH cassette from pBCATPH (Gritz and Davies, 1983). The generated construct was transformed into KJ201 protoplast using PEG-mediated transformation, and transformants were selected on TB3 agar media with 200 µg/mL Hygromycin B. The deletion mutant candidates were screened through mycelial PCR using specific primers **(Supplemental Table 3)**. Δ*rgs1* strains were confirmed by Southern blot analysis and RT-PCR **(Supplemental Fig. 8)**. To produce complementation strains of Δ*rgs1*, we amplified entire gene coding region with 1.5-kb of upstream and downstream from the gDNA of KJ201. We co-transformed this construct with pII99, a vector containing the geneticin-resistant gene, into Δ*rgs1* protoplasts. The complementation strain candidates were selected on TB3 agar medium with 800 µg/mL Geneticin, and confirmed by RT-PCR.

### Phenotype assay

Mycelial growth of wild type and mutant strains was observed after culturing on minimal agar medium (MMA) and modified complete agar medium (CMA) at 25 °C for 9 days. Conidia were collected with 5 mL of sterilized distilled water from cultures on V8 agar medium after incubation for 7 days. Conidiation was measured by counting the number of conidia using a hemocytometer after filtering through one layer of Miracloth (Calbiochem). Conidial germination and appressorium formation were performed on hydrophobic surface. Conidial suspension was adjusted to a concentration of 5 × 10^4^/mL and 70 μL of conidial suspension was dropped on hydrophobic surface with three replicates. After incubation for 2 and 8 h, the rates of conidial germination and appressorium formation was measured by counting at least 100 conidia using Leica DM 750 microscope. To observe conidiophore development, each strain was incubated on oatmeal agar medium for 10 days, and the scraped agar blocks were transferred onto slide glasses. After incubating in the light for 24 h, conidiophore formation was observed under the microscope.

## Author contributions

Y.-J.Y., Y.-J.L., and Y.-H.L. designed the research. Y.-J.Y., Y.-J.L., H.L., S.K., and H.C. performed the experiments and analyzed the data. Y.-J.Y., Y.-J.L., C.H.K., and Y.-H.L. wrote the article. Y.-J.L., and Y.-H.L. acquired research grants.

## Acknowledgement

We thank Prof. Ki-Tae Kim for AlphaFold analysis of DNA-protein interactions.

