## Supplementary information for "*In planta*-specific transcriptional regulatory circuit regulates expression of *MoHTR1*, a nuclear effector gene of *Magnaporthe oryzae*"

**This PDF file includes:**

Supplemental Figures 1 to 7

Supplemental Tables 1 to 3

**Other supporting data for this manuscript include the following:**

Supplemental Data 1 (separate file). List of 415 IPA element carrying genes

Supplemental Data 2 (separate file). Conidial size measurements of WT, $P_{MoHTR1}$:*MobZIP14*, $P_{MobZIP14}$:*MobZIP14*, and *ΔMobzip14*

Supplemental Data 3 (separate file). Transcription factor candidates binding to IPA element selected by expression pattern and pull-down assay

Supplemental Data 4 (separate file). Mass spectrometry results from pull-down assay

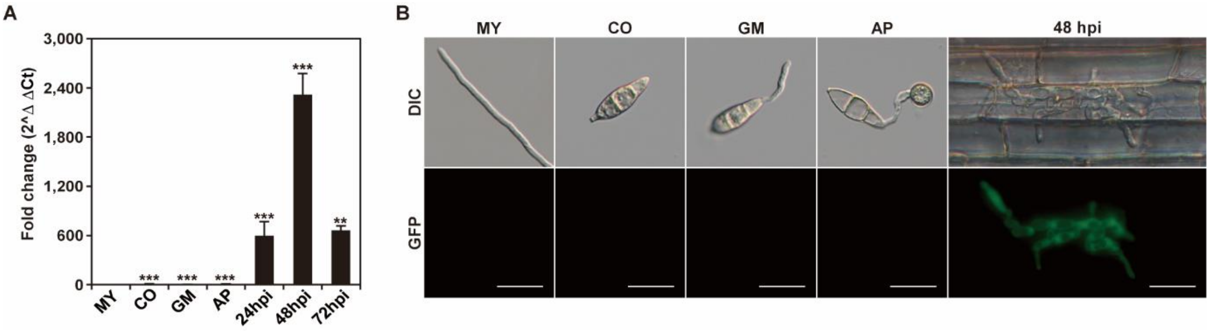


**Supplemental Figure 1. *MoHTR1* is highly expressed during infection stages.**

**(A)** The gene expression pattern of *MoHTR1* at various developmental and infection stages. Relative expression of the gene was measured using quantitative reverse transcription PCR (qRT-PCR). MY, mycelia; CO, conidiation; GM, conidial germination; AP, appressorium formation; Infection stages; hpi, hours of post inoculation. Relative expression was calculated using the $2^{-\Delta\Delta Ct}$ method with mycelial samples as the reference. The *β-tubulin* gene was used for normalization of qRT-PCR data. Mean ± SD, three independent experiments, asterisks indicate statistical significance by an unpaired two-tailed Student's *t*-test ($^{**}p < 0.01$ and $^{***}p < 0.001$). **(B)** GFP fluorescence assay of the *MoHTR1* expression at the various developmental and infection stages. Scale bar indicates 20 μm.

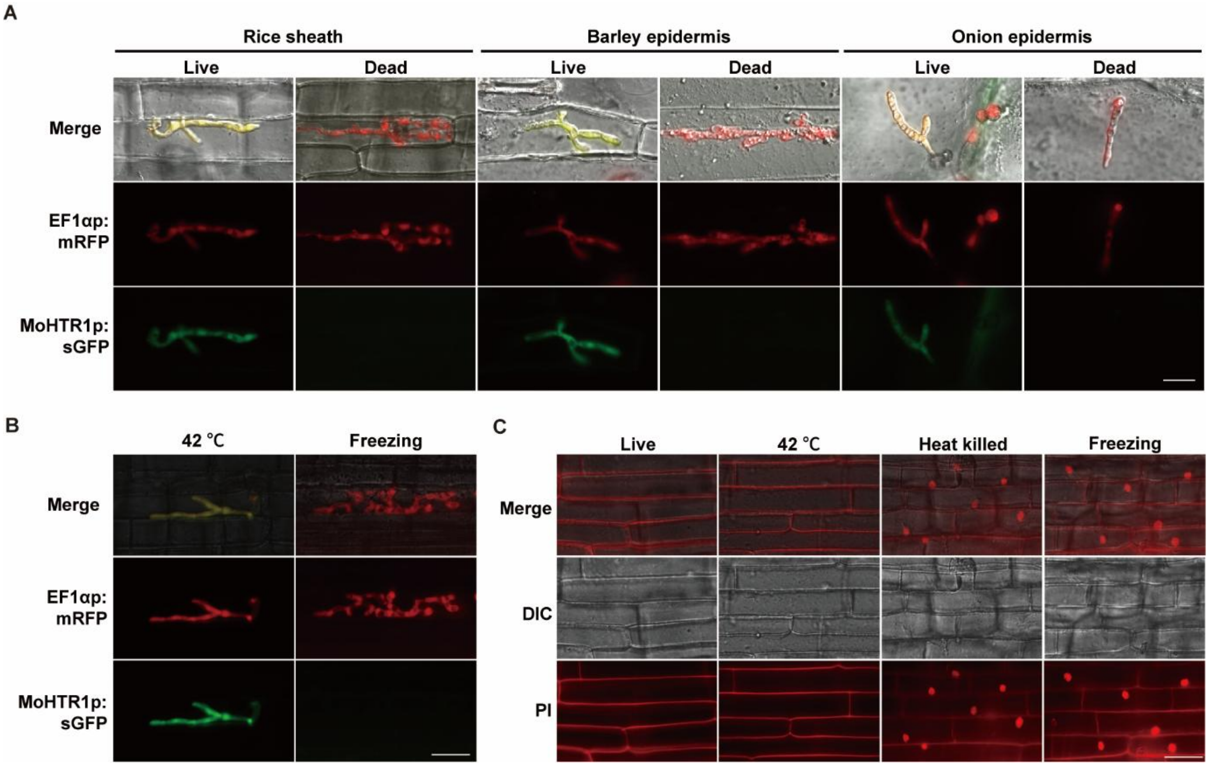


**Supplemental Figure 2. *MoHTR1* is expressed in the living plant cells.**

**(A)** GFP fluorescence images of *MoHTR1* expression in living and dead plant cells. The MoHTR1pro:sGFP construct was co-transformed with EF1αpro:RFP, which serves as a constitutively expressed positive marker. GFP fluorescence was observed at 32 hpi. Scale bar indicates 20 μm. **(B)** Observation of *MoHTR1* expression after treating the heat stress (42 ℃) and freezing using the fluorescence microscope. Scale bar indicates 25 μm. **(C)** To confirm the cell death under each stress condition, we stained rice epidermal cells using 10 μg/mL propidium iodide (PI). Scale bar indicates 25 μm.

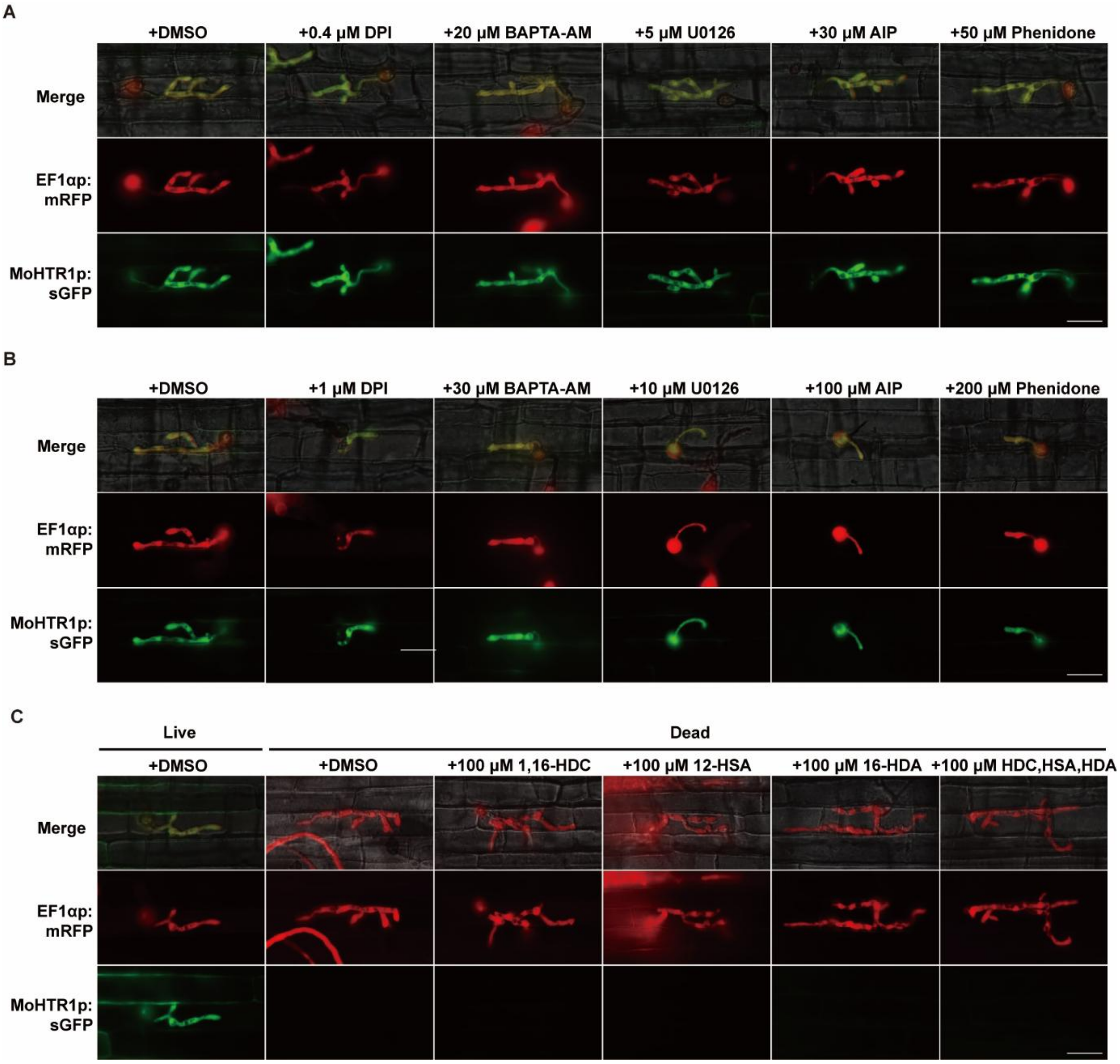


**Supplemental Figure 3.** ***The in planta*-specific expression of MoHTR1 is not regulated by immune signaling pathways and cutin-derived molecules.**

**(A and B)** GFP fluorescence image showing *MoHTR1* expression under treatment of various inhibitors: 0.4 and 1 μM DPI (ROS accumulation inhibitor), 20 and 30 μM BAPTA-AM ($Ca^{2+}$ flux inhibitor), 5 and 10 μM U0126 (MAP kinase cascade inhibitor), 30 and 100 μM AIP (SA signaling pathway inhibitor), and 50 and 200 μM phenidone (JA signaling pathway inhibitor). A fungal transformant co-expressing EF1αpro:RFP and MoHTR1pro:sGFP was inoculated with each inhibitor, and fluorescence signals were observed at 32 hpi. Scale bar indicates 25 μm. **(C)** GFP fluorescence images of *MoHTR1* expression in dead rice sheath cells under treatment of 100 μM cutin monomers. These cutin monomers included 1,16-Hexadecanediol (1,16-HDC), 12-Hydroxystearic acid (12-HSA), and 16-Hydroxyhexadecanoic acid (16-HDA). The fluorescence signals were observed at 32 hpi. Scale bar indicates 25 μm.

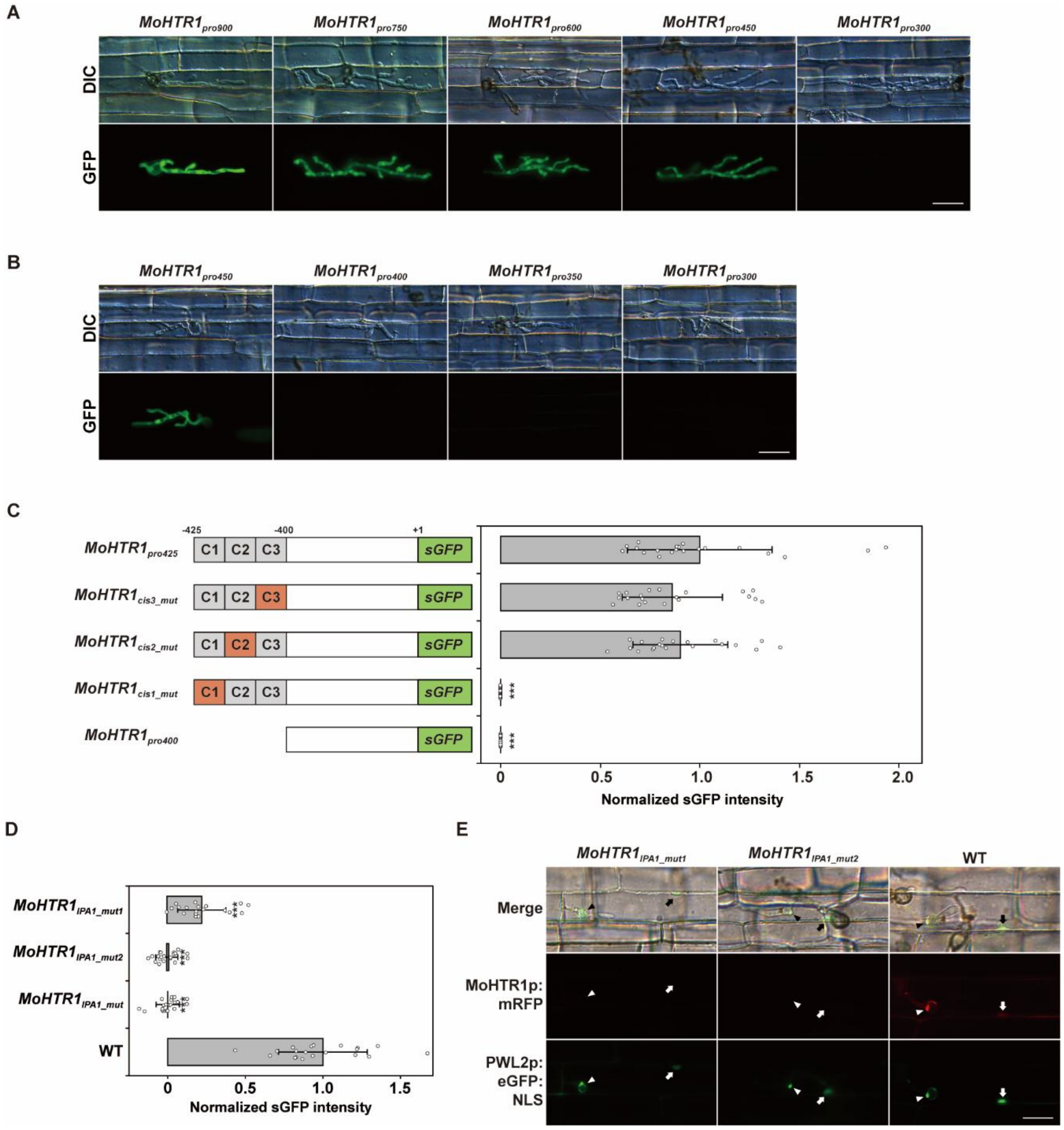


**Supplemental Figure 4. The *in planta*-specific expression of *MoHTR1* is regulated by the IPA element.**

**(A and B)** GFP fluorescence of various truncated *MoHTR1* promoter constructs in the invasive hyphae. GFP signal was detected under a fluorescence microscope at 32 hpi. Scale bar indicates 20 μm. **(C)** Determination of promoter activity of *MoHTR1* promoter constructs fused to GFP by fluorescence intensity. GFP signal was detected under a fluorescence microscope at 32 hpi. To quantify the GFP signal, the fluorescence intensity of invasive hyphae at 20 sites for each strain was measured using image J. Mean ± SD, $n$ =20, three independent experiments, asterisks indicate statistical significance by an unpaired two-tailed Student's *t*-test ($^{***}p < 0.001$). **(D)** Quantification of promoter activity in MoHTR1 IPA element mutants. Each strain was inoculated in rice sheath cells for 32 h. To quantify the GFP signal, the fluorescence intensity of invasive hyphae at 20 sites for each strain was measured using image J. Mean ± SD, $n$ =20, three independent experiments, asterisks indicate statistical significance by an unpaired two-tailed Student's t-test (***$p < 0.001$). **(E)** To observe MoHTR1 localization, we generated fungal transformants expressing MoHTR1 fused to mRFP under the native promoter or under promoters carrying mutations in the IPA element, co-transformed with PWL2:eGFP:NLS, a maker used to label BIC and rice nuclei. Each strain was inoculated in rice sheath cells for 32 h. Scale bar indicates 20 μm.

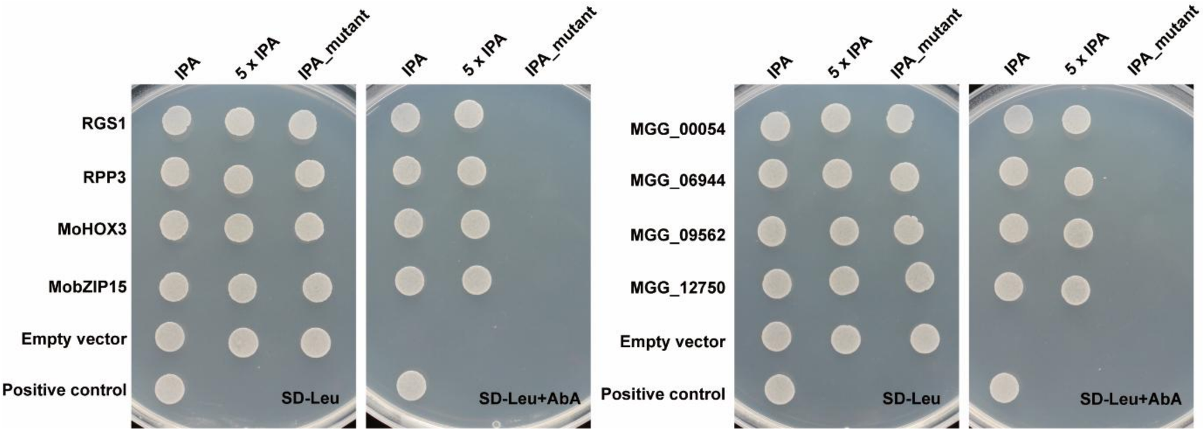


**Supplemental Figure 5. Full plate images of Y1H assays demonstrating transcription factor binding to the IPA element.**

All panels show results from single-plate Y1H experiments in which all controls and test samples were spotted and processed together. The same bait constructs as described in Figure 5A were used: IPA (-442 to -402 region from the start codon of MoHTR1), 5 × IPA (five times repeat sequence of the 8-bp IPA element), and IPA mutant (transversion mutation of the 8-bp of IPA element sequence among 40-bp of promoter regions) inserted into pAbAi vector. Selected transcription factor candidates were cloned into pGADT7 prey vector. pGADT7Rec-p53/p53-AbAi was used as the positive control, and bait-only cells carrying pGADT7 (empty vector, EV) served as the negative control. All constructs were co-transformed into Y1HGold yeast strain and grown on SD/−Leu medium containing AbA (aureobasidin A) for selection.

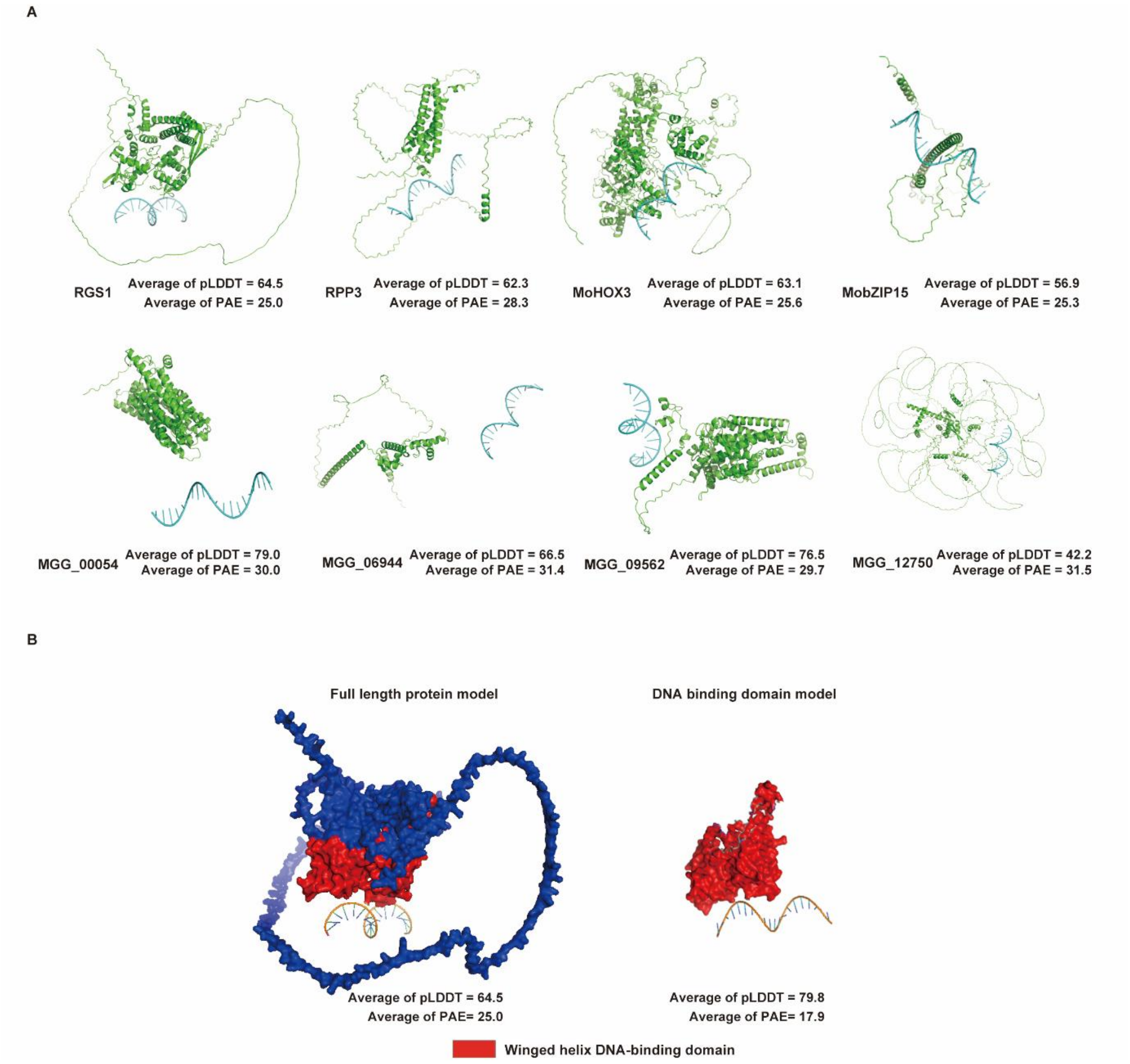


**Supplemental Figure 6. AlphaFold protein–DNA modeling are performed for all eight Y1H-selected TFs with the IPA element.**

**(A)** AlphaFold protein–DNA models of the eight Y1H-selected TFs bound to the IPA probe (full-length protein). Representative 3D views are shown with summarized interface metrics (mean pLDDT and mean PAE) Transcription factor-IPA element complex is predicted by AlphaFold anlaysis. **(B)** Comparison of AlphaFold protein–DNA models for full-length Rgs1 and the isolated DNA-binding domain (DBD) bound to the IPA probe. The DBD model shows higher mean pLDDT and lower inter chain PAE than the full-length model, indicating a more confidently defined binding interface.

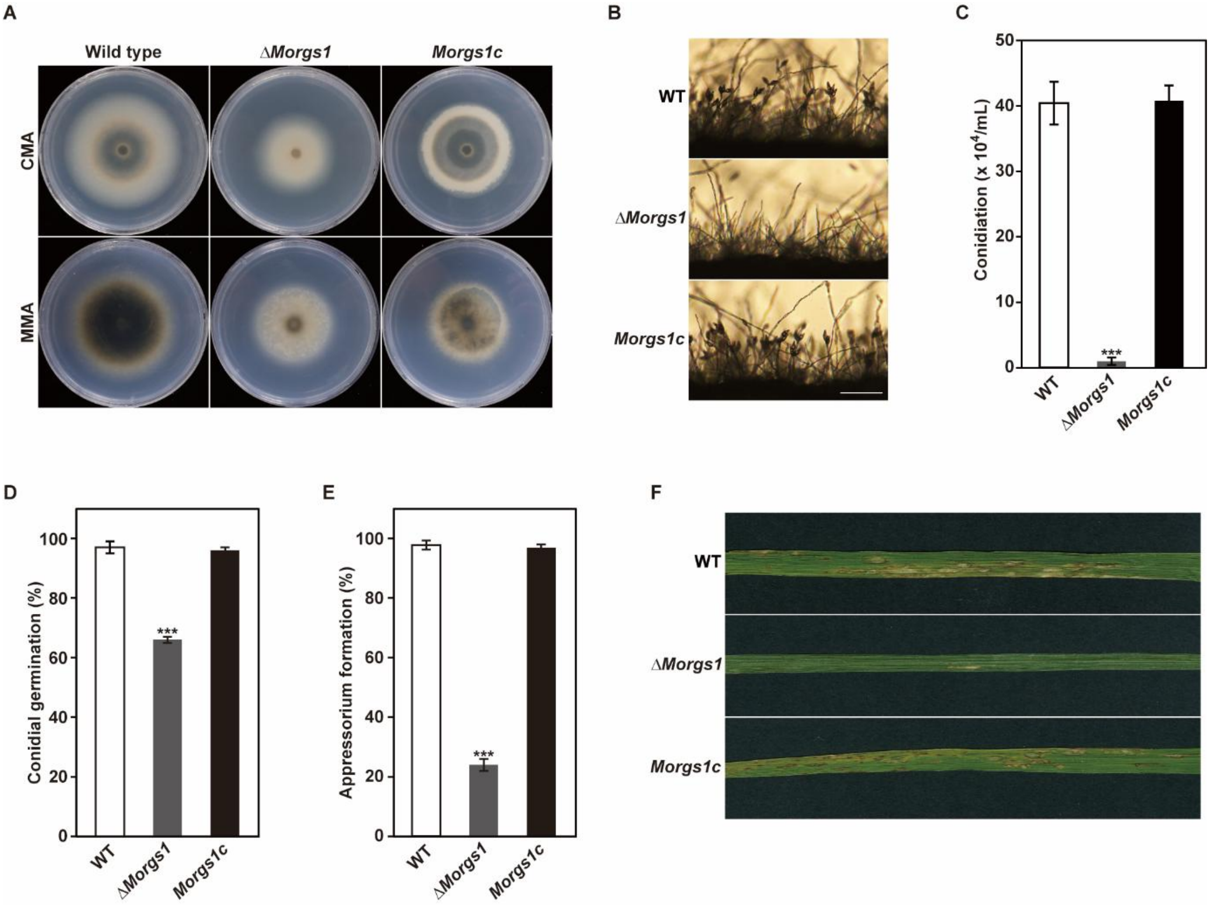


**Supplemental Figure 7. Deletion of *RGS1* affects hyphal growth, conidiation, conidial germination, appressorium formation, and pathogenicity of *M. oryzae***

**(A)** Mycelial growth of the wild type, Δ*Morgs1*, and *Morgs1c* was observed on modified complete agar medium (CMA) and minimal agar medium (MMA) after 9 days of incubation at 25 °C. **(B)** Conidiophore development was observed on oatmeal agar medium under microscope. Scale bar indicates 20 μm. **(C)** Conidia of the wild type, Δ*Morgs1*, and *Morgs1c* were collected from 7-day-old cultures on V8 agar medium. The number of conidia was measured using a hemocytometer under a microscope. Mean ± SD, three independent experiments, significance was determined by an unpaired two-tailed Student's *t*-test ($^{**}p < 0.01$ and $^{***}p < 0.001$). **(D)** Frequency of conidial germination was measured on a hydrophobic surface after 2 h of incubation. Mean ± SD, three independent experiments, significance was determined by an unpaired two-tailed Student's *t*-test ($^{**}p < 0.01$ and $^{***}p < 0.001$). **(E)** Frequency of appressorium formation was measured on a hydrophobic surface 8 h of incubation. Mean ± SD, three independent experiments, significance was determined by an unpaired two-tailed Student's *t*-test ($^{**}p < 0.01$ and $^{***}p < 0.001$). **(F)** To assess pathogenicity, conidial suspensions ($5 \times 10^4$ spores/mL) were inoculated onto 4-week-old Nakdong rice seedlings. Disease lesions were collected at 6 dpi.

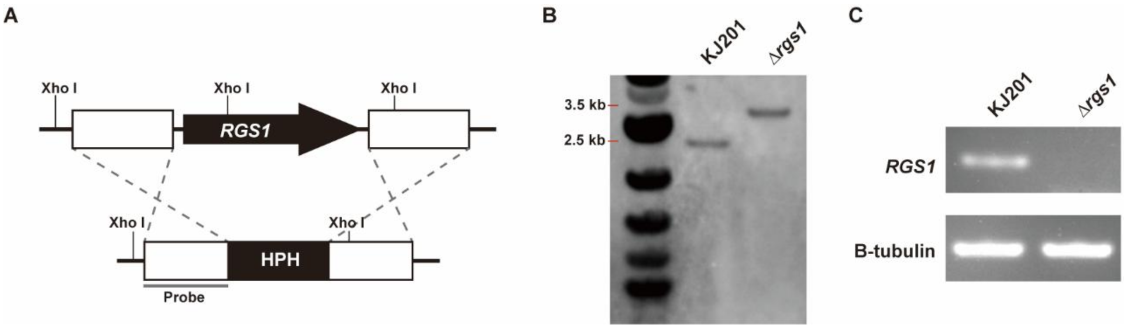


**Supplemental Figure 8. *RGS1* deletion mutant was verified by Southern hybridization and RT-PCR.**

**(A)** Genomic DNA of wild type and Δ*Morgs1* were digested with Xho I. **(B)** Southern blot hybridization of *RGS1* deletion mutant was conducted with upstream flanking region of *RGS1* as a probe. **(C)** RT-PCR analysis was performed to confirm the deletion of *RGS1*. Total RNAs of wild type and Δ*Morgs1* were harvested from mycelium.

**Supplemental Table 1.** qRT–PCR Ct, ΔCt, and ΔΔCt values for the promoter-swapping assays, with statistical significance assessed by two-sided Student’s t-tests.

| **Raw data (Mycelia)** | | Ct | | ΔCt | ΔΔCt | 2^(-ΔΔCt) | Average | STADEV | | | |
|---|---|---|---|---|---|---|---|---|---|---|---|
| | | β-tubulin | MobZIP14 | MobZIP14 | MobZIP14 | MobZIP14 | MobZIP14 | MobZIP14 | | | |
| KJ201 | Rep1 | 15.35 | 18.33 | 2.98 | 0.09 | 0.939522749 | 1.0014629 | 0.066521505 | | | |
| | Rep2 | 15.39 | 18.29 | 2.9 | 0.01 | 0.993092495 | | | | | |
| | Rep3 | 15.36 | 18.15 | 2.79 | -0.1 | 1.071773463 | | | | | |
| $P_{MoHTR1}$ | Rep1 | 15.43 | 19.95 | 4.52 | 1.63 | 0.323088208 | 0.3594793 | 0.032187838 | Student’s t-tests | | |
| | Rep2 | 15.51 | 19.83 | 4.32 | 1.43 | 0.371130893 | | | t | df | P-value (two-tailed) |
| | Rep3 | 15.51 | 19.78 | 4.27 | 1.38 | 0.384218795 | | | 15.0467 | 4 | 0.0001 |
| $P_{MobZIP14}$ | Rep1 | 14.81 | 17.69 | 2.88 | -0.01 | 1.00695555 | 1.016392 | 0.016344425 | Student’s t-tests | | |
| | Rep2 | 14.83 | 17.71 | 2.88 | -0.01 | 1.00695555 | | | t | df | P-value (two-tailed) |
| | Rep3 | 14.78 | 17.62 | 2.84 | -0.05 | 1.035264924 | | | 0.3774 | 4 | 0.725 |

| **Raw data (Infection)** | | Ct | | ΔCt | ΔΔCt | 2^(-ΔΔCt) | Average | STADEV | | | |
|---|---|---|---|---|---|---|---|---|---|---|---|
| | | β-tubulin | MobZIP14 | MobZIP14 | MobZIP14 | MobZIP14 | MobZIP14 | MobZIP14 | | | |
| KJ201 | Rep1 | 23.55 | 27.67 | 4.12 | 0.066666667 | 0.954841604 | 1.0037446 | 0.108105823 | | | |
| | Rep2 | 23.58 | 27.46 | 3.88 | -0.173333333 | 1.127660927 | | | | | |
| | Rep3 | 23.58 | 27.74 | 4.16 | 0.106666667 | 0.92873141 | | | | | |
| $P_{MoHTR1}$ | Rep1 | 25.68 | 26.68 | 1 | -3.053333333 | 8.301277273 | 8.7410323 | 0.423969118 | Student’s t-tests | | |
| | Rep2 | 25.67 | 26.53 | 0.86 | -3.193333333 | 9.147219896 | | | t | df | P-value (two-tailed) |
| | Rep3 | 25.9 | 26.82 | 0.92 | -3.133333333 | 8.774599838 | | | 30.629 | 4 | 0 |
| $P_{MobZIP14}$ | Rep1 | 25.08 | 29.13 | 4.05 | -0.003333333 | 1.002313162 | 0.8192963 | 0.195853826 | Student’s t-tests | | |
| | Rep2 | 25.25 | 29.55 | 4.3 | 0.246666667 | 0.842841545 | | | t | df | P-value (two-tailed) |
| | Rep3 | 25.11 | 29.87 | 4.76 | 0.706666667 | 0.612734221 | | | 1.428 | 4 | 0.226 |

**Supplemental Table 2.** The 45 transcription factor candidates binding to IPA element were selected by pull-down assay.

| Protein name | Locus number | Replicates | TF types | Signal peptide | Reference |
|---|---|---|---|---|---|
| MoCOD2 | MGG_09263 | 3 | Zn2Cys6 | X | (Chung *et al.*, 2013) |
| MoHTR4 | MGG_05518 | 2 | C2H2 zinc finger | O | (Kim *et al.*, 2020) |
| MoHOX3 | MGG_01730 | 2 | Homeobox | X | (Kim *et al.*, 2009) |
| RGS1 | MGG_14517 | 2 | Winged helix repressor DNA-binding | X | (Zhang *et al.*, 2011) |
| MoHOX6 | MGG_11712 | 2 | Homeobox | X | (Kim *et al.*, 2009) |
| - | MGG_06944 | 2 | C2H2 zinc finger | X | - |
| - | MGG_09562 | 2 | Zn2Cys6 | X | - |
| - | MGG_00054 | 2 | Myb | X | - |
| - | MGG_11160 | 2 | Zn2Cys6 | X | - |
| MoFOX2 | MGG_14496 | 1 | Forkhead | X | (Kim *et al.*, 2009) |
| MoMyb12 | MGG_01130 | 1 | Myb | X | (Li et al., 2021) |
| MoHOX7 | MGG_12865 | 1 | Homeobox | X | (Kim *et al.*, 2009) |
| MIG1 | MGG_01204 | 1 | MADS-box | X | (Mehrabi *et al.*, 2008) |
| MoHOX2 | MGG_00184 | 1 | Homeobox | X | (Kim *et al.*, 2009) |
| RPP5 | MGG_08917 | 1 | Zn2Cys6 | X | (Osés-Ruiz *et al.*, 2021) |
| RPP3 | MGG_07218 | 1 | Zn2Cys6 | X | (Osés-Ruiz *et al.*, 2021) |
| CNF3 | MGG_13360 | 1 | Zn2Cys6 | X | (Lu et al., 2014) |
| MoHTR3 | MGG_04580 | 1 | C2H2 zinc finger | O | (Lee et al., 2023) |
| MoHTR15 | MGG_10494 | 1 | Myb | X | (Kim *et al.*, 2020) |
| - | MGG_05239 | 1 | Winged helix repressor DNA-binding | X | - |
| - | MGG_08340 | 1 | C2H2 zinc finger | X | - |
| - | MGG_02732 | 1 | bZIP | X | - |
| - | MGG_06529 | 1 | C2H2 zinc finger | X | - |
| - | MGG_06832 | 1 | Zn2Cys6 | X | - |
| - | MGG_06258 | 1 | Forkhead | X | - |
| - | MGG_00139 | 1 | C2H2 zinc finger | X | - |
| - | MGG_09837 | 1 | C2H2 zinc finger | X | - |
| - | MGG_00076 | 1 | C2H2 zinc finger | X | - |
| - | MGG_03538 | 1 | GATA type zinc finger | X | - |
| - | MGG_04521 | 1 | GATA type zinc finger | X | - |
| - | MGG_05035 | 1 | bZIP | X | - |
| - | MGG_04665 | 1 | Zn2Cys6 | X | - |
| - | MGG_01853 | 1 | Forkhead | X | - |

| | | | | | |
|---|---|---|---|---|---|
| - | MGG_14070 | 1 | C2H2 zinc finger | X | - |
| - | MGG_09780 | 1 | Zn2Cys6 | X | - |
| - | MGG_07699 | 1 | C2H2 zinc finger | O | - |
| - | MGG_06090 | 1 | C2H2 zinc finger | X | - |
| - | MGG_00702 | 1 | C2H2 zinc finger | X | - |
| - | MGG_15357 | 1 | Myb | X | - |
| - | MGG_13986 | 1 | Homeobox | X | - |
| - | MGG_05615 | 1 | Winged helix repressor DNA-binding | X | - |
| - | MGG_01946 | 1 | Zn2Cys6 | X | - |
| - | MGG_10162 | 1 | C2H2 zinc finger | X | - |
| - | MGG_07319 | 1 | GATA type zinc finger | X | - |

**Supplemental Table 3.** Primers used in this study.

| **Primer name** | **Sequences (5' – 3')** |
|---|---|
| **Primers for promoter truncation** | |
| MoHTR1_900pro_SpeI_F | ACTAGTGCTTACAAAGTCTAATTGATATGCATG |
| MoHTR1_750pro_SpeI_F | ACTAGTTAGAATGGGCACTGCCGG |
| MoHTR1_600pro_SpeI_F | ACTAGTATCGAGGCTTTGTTGTCATACTG |
| MoHTR1_450pro_SpeI_F | ACTAGTTAATACTATTCATGAAAGTCAATCATATTTCG |
| MoHTR1_425pro_SpeI_F | ACTAGTTATTTCGTGTACAGATTTGGATG |
| MoHTR1_400pro_SpeI_F | ACTAGTGACTGTTTCCGCAAATGG |
| MoHTR1_350pro_SpeI_F | ACTAGTAATAGACATTAATGCACACCAGG |
| MoHTR1_300pro_SpeI_F | ACTAGTATACTATTGCCATATTTACCAAGACAAG |
| MoHTR1_IPA1_mut_SpeI_F | ACTAGTGCGGGATGTGCAGATTTGGATGTTGACTGTTTCC |
| MoHTR1_IPA2_mut_SpeI_F | ACTAGTTATTTCTGTGCACTCGGTGGATGTTGACTGTTTCCG |
| MoHTR1_IPA3_mut_SpeI_F | ACTAGTTATTTCGTGTACATCGGGTTCGTGGTTGACTGTTTCCGC |
| MoHTR1_pro_NcoI_R | CCATGGTTTGGCAGAGGTATGTCACG |
| sGFP_NcoI_F | CCATGGATGGTGAGCAAGGGCGA |
| sGFP_ApaI_R | GGGCCCTTACTTGTACAGCTCGTCCATGC |
| **Primers for IPA element carrying genes** | |
| Slp1_qRT_F | GATGTTGTCGATGACGCAGG |
| Slp1_qRT_R | ATCGTGTCGCAGAAGCTCAA |
| MoHDA1_qRT_F | CTGACGGTGGTAACTGTGCT |
| MoHDA1_qRT_R | CCCCGTGTCCTTCGAATAGG |
| MGG_08940_qRT_F | CTCCCGTACGAAGCCATTGT |
| MGG_08940_qRT_R | TTACCCTCTTGTCGCTGACG |
| MGG_15336_qRT_F | GCAGATATGGGTAAGTGTAACTGTG |
| MGG_15336_qRT_R | TAACGCCGTCCGTAATGGTG |
| MGG_18035_qRT_F | TAGCCCAATATGCACTCGCC |
| MGG_18035_qRT_R | GCAAAGGAAAGGTTTCGTGCT |
| MGG_09666_qRT_F | GGTCAGCTACGTCAACCACA |
| MGG_09666_qRT_R | TTCACGGTAGTTTCGGGCTG |
| Slp1_pro_SpeI_F | ACTAGTACCAAGGTTGAGTAGAAAACGA |
| Slp1_IPA_mut_ SpeI _F | ACTAGTACCAAGGTTGAGTAGAAACATCCCGCTTG |
| Slp1_pro_ EcoRI_R | GAATTCTTGACGGTTTGAGAGACGG |
| sGFP_EcoRI_F | GAATTCATGGTGAGCAAGGGCGA |
| **Primers for promoter switching** | |
| MoHTR1_pro_ApaI_F | GGGCCCTATTTCGTGTACAGATTTGG |
| MoHTR1_pro_SpeI_R | ACTAGTTTTGGCAGAGGTATGTCACG |
| MobZIP14_gene_SpeI_F | ACTAGTCTTGTGCAGACTCCACAC |
| MobZIP14_gene_NcoI_R | CCATGGCGCTTGGAAGTCTGCG |
| MobZIP14_qRT_F | GGATAGTCGCCTTGTGTCGT |
| MobZIP14_qRT_R | AGAGCAACAGGTGCAAGAGC |
| **Primers for pull-down assay** | |
| IPA probe_sense | GAGCTGATGGGATGTCGAAGCTAATACTATTCATGAAAGTCAATCATATTTCGTGTACAGATTTGGATGTTGACTGTTTCCGCAAATGGAAAGATTGTCA |
| IPA probe_antisense | TGACAATCTTTCCATTTGCGGAAACAGTCAACATCCAAATCTGTACACGAAATATGATTGACTTTCATGAATAGTATTAGCTTCGACATCCCATCAGCTC |
| IPA mutant probe_sense | GAGCTGATGGGATGTCGAAGCTAATACTATTCATGAAAGTCAATCAGCGGGATGGTACAGATTTGGATGTTGACTGTTTCCGCAAATGGAAAGATTGTCA |
| IPA mutant probe_antisense | TGACAATCTTTCCATTTGCGGAAACAGTCAACATCCAAATCTGTACCATCCCGCTGATTGACTTTCATGAATAGTATTAGCTTCGACATCCCATCAGCTC |

**Supplemental Table 3.** Continued

| **Primers for yeast one hybrid** | |
|---|---|
| MGG_13894_NdeI_F | CATATGATGGAACCCACAAATGATGGGA |
| MGG_13894_ClaI_R | ATCGATCTACCTCAACGACGGTAGCC |
| MGG_12776_NdeI_F | CATATGATGGCGGAGAGAGTAGAAAGAC |
| MGG_12776_ClaI_R | ATCGATCTAGTTTGGCACACATGCAGG |
| MGG_04699_NdeI_F | CATATGATGACGATGACGTTAGACAGCAAC |
| MGG_04699_ClaI_R | ATCGATTTAGTCAGAGTGATGGTCCTCGG |
| MGG_12750_SmaI_F | CCCGGGATGTCATCAACCAACGGCTTGG |
| MGG_12750_ClaI_R | ATCGATTTATGTACGGGCAGCGAACG |
| MGG_07305_NdeI_F | CATATGATGCTCGCAAGGAGGGG |
| MGG_07305_ClaI_R | ATCGATTTAGATGTGGCCATAGCTCGTAAT |
| MGG_05369_NdeI_F | CATATGATGCCCGCTTTCTTTGACCC |
| MGG_05369_ClaI_R | ATCGATTCAACCACTCTTCCTTGGACG |
| MGG_09780_NdeI_F | CATATGATGCCTTATACGTGCGAAGTCG |
| MGG_09890_ClaI_R | ATCGATTTAGGCTCCGAGAGATAACATGATTTCATC |
| MGG_01207_NdeI_F | CATATGATGACGGAAAAGGATGCTGCC |
| MGG_01207_ClaI_R | ATCGATTTACGCATCGTTGAATCCACCG |
| MoCOD2_SmaI_F | CCCGGGATGGACCACGGCCCGGACCCTAAGCGG |
| MoCDC2_XhoI_R | CTCGAGTCATCGAGGTACCTGAACATCCACTG |
| MoHOX3_SmaI_F | CCCGGGATGCCAATAGTACCAGAAGAAGACG |
| MoHOX3_ClaI_R | ATCGATTTACATGTCGAAAAGCTGACTTGG |
| RGS1_NdeI_F | CATATGATGGACGACACCTCCCG |
| RGS1_EcoRI_R | GAATTCCTATAACCGTTGCGAGCGGC |
| MoHOX6_NdeI_F | CATATGATGGCCTTCCCTTCCCAG |
| MoHOX6_EcoRI_R | GAATTCTTATCGCCAGTTGCCCTTTG |
| MGG_06944_NdeI_F | CATATGATGGGGAAAGCAGAGTTCGG |
| MGG_06944_EcoRI_R | GAATTCTTACATCCTTTGGCGTTTACTAGGG |
| MGG_09562_NdeI_F | CATATGATGGTCTACTGCGGAAAGCC |
| MGG_09562_EcoRI_R | GAATTCTTAGACAGCCAGCATCTTCTGGG |
| MGG_00054_EcoRI_F | GAATTCATGGCCGATATCGAAAAAGTCACAC |
| MGG_00054_SacI_R | GAATTCTCACAAAATATAGCGGAACTGCTTG |
| MGG_11160_NdeI_F | CATATGATGAGCTACATACGAAAACGCG |
| MGG_11160_EcoRI_R | GAATTCTTAGTCCCTTGTTGTAATCCGCTC |
| MoHOX7_NdeI_F | CATATGATGGAATATACCTTGCCAAACCACC |
| MoHOX7_EcoRI_R | GAATTCTTAGACGCTGCCACGCT |
| MIG1_NdeI_F | CATATGATGGGTCGCAGGAAGATTGA |
| MIG1_EcoRI_R | GAATTCTTAAGAGTCTACTTTGACCCGTTTTGG |
| MoHOX2_ClaI_F | ATCGATATGGACTACATGAACATGTTTCAGG |
| MoHOX2_SacI_R | GAGCTCTTACTTCTCGACACCTGTGTTGTAG |
| RPP3_EcoRI_F | GAATTCATGTCAGCCTCTGCAAAGCC |
| RPP3_SacI_R | GAGCTCTTATACAAACCCCATATTATCCTTTCGGTG |
| CNF3_NdeI_F | CATATGATGAGAGAGTTACTATCGTGTTCAGTCTGTAGGC |
| CNF3_EcoRI_R | GAATTCTCATAATTCATTAGCTATCCGGTCACACGTGGC |
| MoHTR15_NdeI_F | CATATGATGTGGACCTTCTTAATACTCTGCGCAATTTTGTTCTGCG |
| MoHTR15_EcoRI_R | GAATTCTCATTCGACAAGACCCAGAATACGCGCGACC |
| IPA_bait_SacI/XhoI_F | GAGCTCTCATGAAAGTCAATCATATTTCGTGTACAGATTTGGATGT CTCGAG |
| IPA_bait_SacI/XhoI_R | CTCGAGACATCCAAATCTGTACACGAAATATGATTGACTTTCATGA GAGCTC |
| 5 x IPA_bait_SacI/XhoI_F | GAGCTCTATTTCGTTATTTCGTTATTTCGTTATTTCGTTATTTCGTCT CGAG |
| 5 x IPA_bait_SacI/XhoI_R | CTCGAGACGAAATAACGAAATAACGAAATAACGAAATAACGAAATA GAGCTC |
| IPA_mutant_SacI/XhoI_F | GAGCTCTCATGAAAGTCAATCAGCGGGATGGTACAGATTTGGATG TCTCGAG |

**Supplemental Table 3.** Continued

| | |
|---|---|
| **Primers for yeast one hybrid** | |
| IPA_mutant_SacI/XhoI_R | CTCGAGACATCCAAATCTGTACCATCCCGCTGATTGACTTTCATGA GAGCTC |
| **Primers for electrophoretic mobility shift assay** | |
| RGS1_EcoRI_F | GAATTCATGGACGACACCTCCCG |
| RGS1_HindIII_R | AAGCTTCTATAACCGTTGCGAGCGGC |
| **Primers for qRT-PCR analysis of IPA element-containing genes** | |
| RGS1_qRT_F | AAAGCGCTGAAGCATTGACC |
| RGS1_qRT_R | CGCTCAAGTTCTCCCAGTCA |
| MoHTR1_qRT_F | GCCCTGGAATCAATACGGAG |
| MoHTR1_qRT_R | GAGTCTATCGTTGCCAGGAAG |
| Slp1_qRT_F | GATGTTGTCGATGACGCAGG |
| Slp1_qRT_R | ATCGTGTCGCAGAAGCTCAA |
